# Proximity proteomics reveals UCH-L1 as an NLRP3 interactor that modulates IL-1β production in human macrophages and microglia

**DOI:** 10.1101/2023.10.09.561576

**Authors:** Zhu Liang, Andreas Damianou, Iolanda Vendrell, Edward Jenkins, Frederik H. Lassen, Sam J Washer, Guihai Liu, Gangshun Yi, Hantao Lou, Fangyuan Cao, Xiaonan Zheng, Ricardo A. Fernandes, Tao Dong, Edward W. Tate, Elena Di Daniel, Benedikt M Kessler

## Abstract

Activation of the NACHT, LRR family pyrin domain containing 3 (NLRP3) inflammasome complex is an essential innate immune signalling mechanism. To reveal how human NLRP3 inflammasome assembly and activation are controlled, in particular by components of the ubiquitin system, proximity labelling, affinity purification and RNAi screening approaches were performed. Our study provides an intricate time-resolved molecular map of different phases of NLRP3 inflammasome activation. Also, we show that ubiquitin C-terminal hydrolase 1 (UCH-L1) interacts with the NACHT domain of NLRP3. Downregulation of UCH-L1 decreases pro-IL-1β levels. UCH-L1 chemical inhibition with small molecules interfered with NLRP3 puncta formation and ASC oligomerization, leading to altered IL-1β cleavage and secretion, particularly in microglia cells, which exhibited elevated UCH-L1 expression as compared to monocytes/macrophages. Altogether, we profiled NLRP3 inflammasome activation dynamics and highlight UCH-L1 as an important modulator of NLRP3-mediated IL-1β production, suggesting that a pharmacological inhibitor of UCH-L1 may decrease inflammation-associated pathologies.

## INTRODUCTION

Innate immunity reflects a cornerstone of defence systems in living organisms which integrates with adaptive immune responses in the case of higher eukaryotes (Paludan et al., 2021). A major hallmark is the activation of inflammasome complexes, composed of ASCs and NLRs, such as NACHT, LRR family pyrin domain containing 3 (NLRP3) (Martinon et al., 2002) involving cleavage of pro-IL1β by caspase-1 (Gross et al., 2012). External stimuli including lipopolysaccharide (LPS) as the first, and nigericin, monosodium urate (MSU) monohydrate, Alum, ATP or cholesterol crystals as the second signal lead to the assembly and activation of the NLRP3 inflammasome complex, resulting in inflammatory responses (Shao et al., 2015; Swanson et al., 2019). Assembly of this supramolecular structure, featuring NLRP3 and ASC, is further regulated by associated factors, such as NEK7 (He et al., 2016), as well as specific posttranslational modifications including phosphorylation and ubiquitylation (Akther et al., 2021; Liang et al., 2021; Zangiabadi and Abdul-Sater, 2022). In addition to the kinases RIPK3 (Lawlor et al., 2015), FYN (Panicker et al., 2019) and ubiquitin ligases TRIM31(Song et al., 2016), gp78/Insig-1 (Xu et al., 2022), MARCH7 (Yan et al., 2015), HUWE1 (Guo et al., 2020), RNF25 and Cbl (Tang et al., 2020), deubiquitylases (DUBs) have been shown to modulate ubiquitylation of components of the inflammasome often in the context of specific diseases (Lopez-Castejon, 2020; Lopez-Castejon and Edelmann, 2016; Lopez-Castejon et al., 2013). More specifically, ubiquitylation and deubiquitylation of NLRP3 and IL-1β (Vijayaraj et al., 2021) have been reported to reflect critical checkpoints of NLRP3 inflammasome complex assembly, activation and signalling (Lopez-Castejon, 2020; Lopez-Castejon and Edelmann, 2016; Lopez-Castejon et al., 2013). Ubiquitin protease 7 (USP7) and 47 (USP47) inhibition resulted in reduced inflammasome activation (Palazon-Riquelme et al., 2018). In addition, in mice, BRCC3, a DUB reflecting a component of the BRISC complex, was shown to deubiquitylate NLRP3, thereby regulating inflammasome activity (Py et al., 2013). Recently, increasing number of findings suggest a complex interplay between ubiquitylation and deubiquitylation mechanisms that modify components of the inflammasome complex. However, most studies of ubiquitin-mediated NLRP3 pathway were conducted in murine models, such as NG5, mouse peritoneal macrophages, and mouse bone marrow-derived macrophages (BMDMs) (Py et al., 2013; Ren et al., 2019; Song et al., 2020). Given the inter-species divergence between human and mouse inflammasome complexes and regulatory pathways, the role of ubiquitylation/deubiquitylation in human NLRP3 inflammasome activation remains largely unknown.

To gain a more comprehensive understanding of the ubiquitin-mediated regulation of the human NLRP3 inflammasome, we have combined ascorbic acid peroxidase 2 (APEX2)-based proximity labelling, affinity purification and RNAi screening to discover molecular factors involved in the primary and secondary signals of NLRP3 inflammasome activation and assembly. In both, a reconstituted system and macrophages, we demonstrate that UCH-L1 interacts with the NACHT domain of NLRP3 and thereby interferes with ASC and NLRP3 assembly. Furthermore, ubiquitin C-terminal hydrolase 1 (UCH-L1) knockdown or chemical inhibition interferes with IL-1β production, particularly in microglia cells that exhibited elevated UCH-L1 expression as compared to peripheral monocytes or macrophages.

## RESULTS

### NLRP3-APEX2 captures the NLRP3 molecular proximity network

As the N-terminus NLRP3 PYD domain has been reported to be responsible for initiating NLRP3 oligomerization, we inserted APEX2 at the C-terminus (LRR domain) of NLRP3 including a flexible linker between these domains and a flag epitope at the N-terminus (**Figure 1A**). To avoid unstable expression and mis-localization during transient transfection, we generated a NLRP3-APEX2 HEK293 stable cell line using the Flp-In T-REx system, allowing dose-dependent control of expression with tetracycline (**Figures S1A, B**). Fluorescence microscopy confirmed that both flag-NLRP3 and flag- NLRP3-APEX2 can form puncta in the perinuclear areas in response to nigericin stimulation, while the fusion proteins are diffused across the cytosol in the resting state (**Figures S1C, D**). ASC oligomerization assays with co-expression of adaptor protein ASC in stable cell lines proved that both flag-NLRP3 and flag-NLRP3-APEX2 are competent to recruit ASC and induce its polymerization upon stimulation with nigericin (**Figures S2A, B**). Taken together, these results indicate that APEX2 tagging fused to the C-terminus of NLRP3 does not impede its ability to sense stimuli, particularly nigericin induced potassium efflux, and recruit ASC to form oligomers.

**Figure 1.**
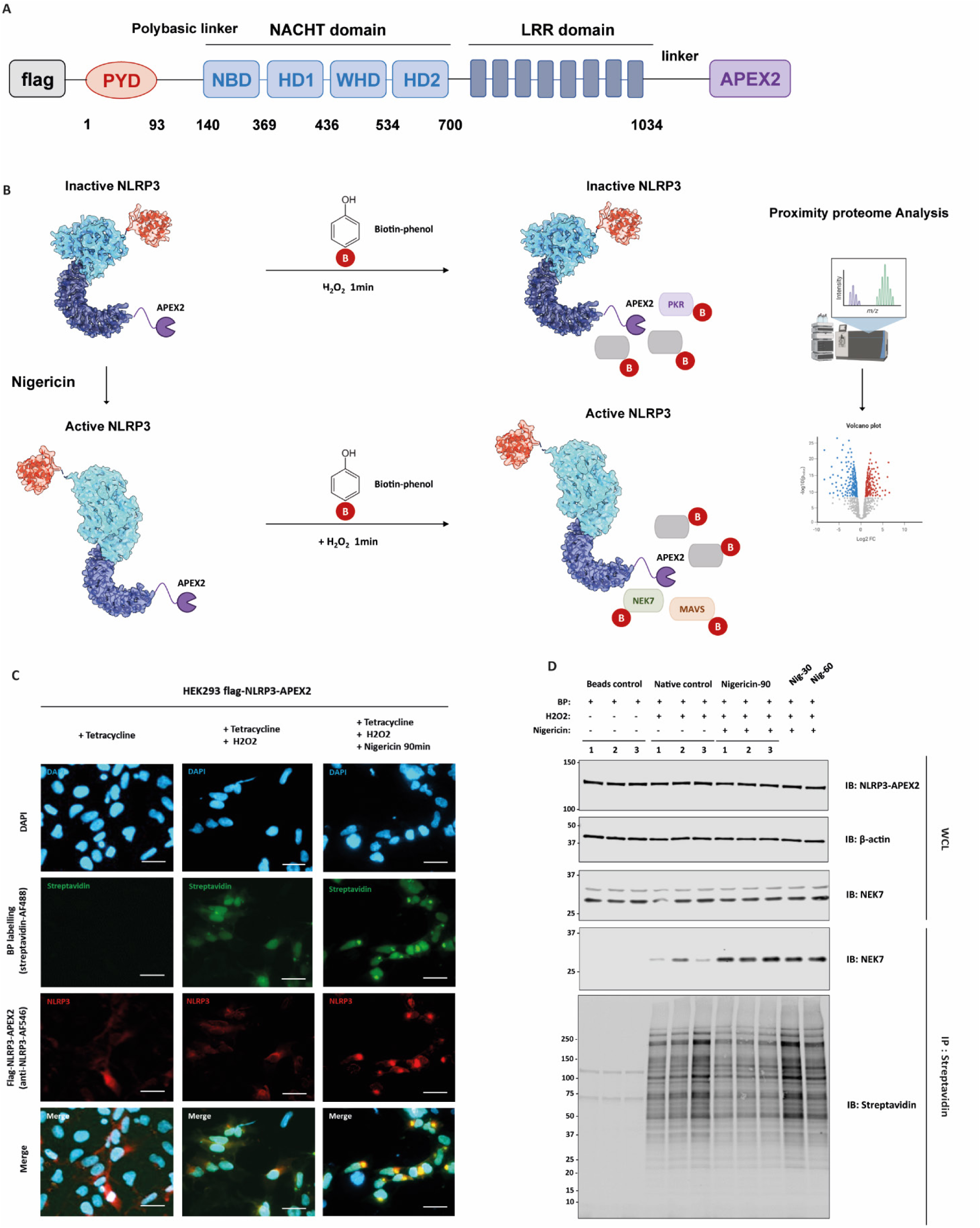
Proximity proteomic profiling of the NLRP3 inflammasome upon activation. (A) Scheme of APEX2-tagging NLRP3. APEX2 was fused to the C terminus of full-length NLRP3 via a glycine-serine linker. (B) Proximity labelling and mass spectrometric analysis workflow for inactive (PDB: 6NPY) and active NLRP3 (PDB: 8EJ4) (Sharif et al., 2019; Xiao et al., 2023). (C) Confocal fluorescence imaging of NLRP3 APEX2 labelling in HEK293 cell lines. NLRP3-APEX2 HEK293 stable cell lines were treated with tetracycline overnight, followed by the treatment nigericin (10μM) or DMSO (control) for 90 min. Subsequently, cells were incubated with biotin phenol, followed by H_2_O_2_ as indicated. Afterwards, cells were fixed and stained with streptavidin-Alexa Fluor 488 (AF488) conjugate to visualize biotinylated proteins and anti-NLRP3 antibody to visualize the localization of NLRP3-APEX2. BP, biotinylated proteins. Scale bars, 50 μm. (D) Immunoblot showing the biotinylated NEK7 captured by streptavidin beads in negative control, unstimulated and nigericin-stimulated HEK293 Flp-In T-REx NLRP3-APEX2 cells. Cells were treated with tetracycline overnight, and stimulated with nigericin for 30 min, 60 min, and 90 min before the APEX2 labelling. Whole cell lysates and streptavidin pull-down were collected and blotted for NLRP3 and NEK7.

We next investigated whether biotin labelling profiles are spatially restricted to the fusion bait. Live HEK293 cells expressing NLRP3-APEX2 were treated with or without nigericin. Subsequently, cells were treated with H_2_O_2_ in the presence of biotin-phenol, followed by rapid quenching, methanol fixation and staining (**Figure 1B**). In contrast to the native control without H_2_O_2_ treatment, we observed robust biotinylation by streptavidin-Alexa Fluor 488 imaging. Under unstimulated conditions, NLRP3-APEX2 was diffused across the cytosol, with biotinylated proteins following similar distribution pattern. Upon nigericin treatment, NLRP3-APEX2 oligomerized to form puncta close to the nucleus, and we also observed specks formed by biotin labelled proteins (**Figure 1C**). This partially suggests the spatial specificity of NLRP3-APEX2 induced biotinylation.

The ability of APEX2 to efficiently label and capture the NLPR3 “proximitome” was assessed using streptavidin immunoblotting and Coomassie blue staining, confirming the robust biotin labelling activity of APEX2 fused to NLRP3 (**Figure 1D, Figure S3B**). Initially, we investigated whether APEX2 is competent to capture the known NLRP3-interacting protein-NIMA-related kinase 7 (NEK7). In contrast to the unlabelled negative control, we found that NLRP3-APEX2 specifically biotinylated NEK7. Additionally, a substantial increase in NEK7 protein sequestration was observed in association with activated NLRP3 (**Figure 1D**).

Given the rapid kinetic properties of APEX2 proximity labelling, we next sought to map the spatiotemporally resolved “proximitome” of NLRP3 upon stimulation by nigericin across various time intervals. Following mass spectrometric analysis, 3,478 proteins in total were detected across the DMSO control and nigericin treatment for 30 min, 60 min and 90 min. To maximize the signal-to-noise ratio and discriminate the bona fide interacting partners from the background contaminants, Bayesian model-based algorithm SAINTexpress (Teo et al., 2014) was applied, yielding 800-1,100 high confidence NLRP3 proximal proteins (**Figures 2A-B, Figure S3A**). Unsupervised principal component analysis (PCA) showed the clustering and patterns of different protein subgroups (**Figure 2C**). High correlation was observed between the three biological repeats within each group confirming the low technical variation in this experiment (**Figure S3C**).

**Figure 2.**
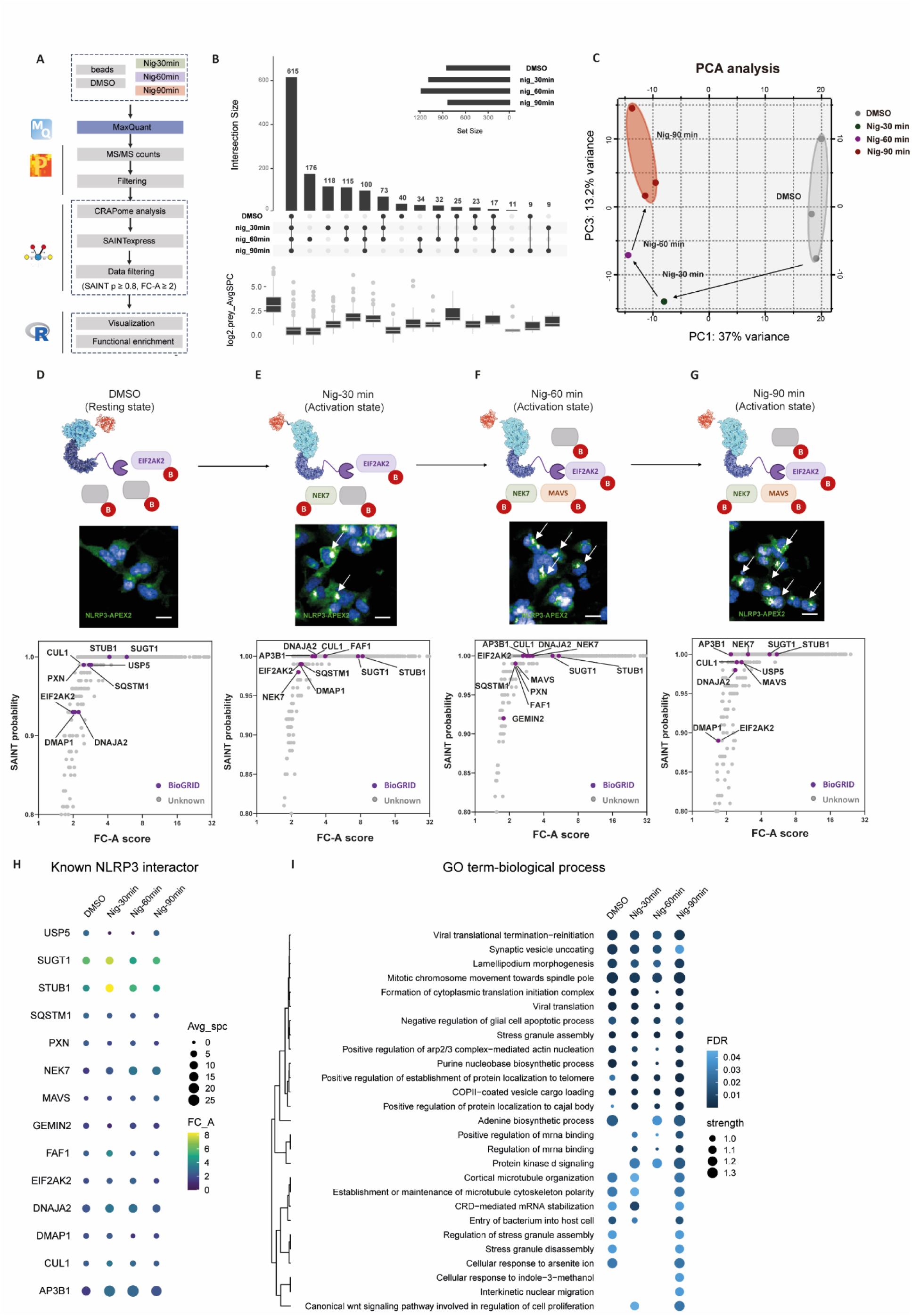
Temporal tracking of the NLRP3 proximity proteome. (A) Schematic illustration of the data analysis for APEX2 PL-MS. (B) The upset plot depicting the intersections of high confidence NLRP3 proximal proteins across samples. The bars in the y axis represent the number of high confidence proximal proteins (SAINT probability score ≥ 0.8, fold change ≥ 2) identified in each sample. The boxplot represents the log2 (average of spectral counts) of the proximal proteins identified in each group. The median, first and third quartiles of the data were displayed as the centre line, the lower and upper boundaries. (C) Principal component analysis of the 8 samples (3 unstimulated DMSO controls, 3 stimulated samples with 90 min treatment of nigericin, 30min nigericin-stimulated samples, and 60 min nigericin-stimulated samples). Each dot represents an individual samples. (D-G) SAINT probability scoring of the proteomic data for NLRP3-APEX2 compared to the negative beads control. Proteins passing the cut off (SAINT probability score ≥ 0.8 and FC-A score ≥ 1) are considered as high confidence proximal proteins for NLRP3. Previously reported NLRP3-interacting proteins (BioGRID, v4.4.213) are coloured in purple. All other proteins are shown in grey. Inactive (PDB: 6NPY) and active NLRP3 (PDB: 8EJ4) are shown as surface view with model. (H) Dot plot of previously reported NLRP3-interacting proteins that are detected in the NLRP3-APEX2 proximitome upon nigericin stimulation for different time points (unstimulated, 30 min, 60 min, and 90 min). Sizes denote the average spectral counts and colours denote the FC-A score. (I) Dot plots displaying the GOBP enrichment analysis of high confidence NLRP3 proximal proteins upon nigericin stimulation for different timepoints as indicated.

To assess the specificity of the NLRP3 proximity proteome, we next sought to check if the known NLRP3 interactors or regulators are biotinylated and captured by APEX2. SAINT probability scoring and simpler fold-change (FC) were calculated by the comparison of prey protein abundance between the bait and negative beads control. Previously reported 83 human NLRP3 interactors, retrieved from public interaction repository BioGrid (https://thebiogrid.org/, version 4.4.209) and the literature (**Table S1**) (Oughtred et al., 2021; Oughtred et al., 2019; Stark et al., 2006) were mapped and plotted with stringent thresholds (SAINT probability > 0.8, FC-A ≥ 1). After filtering, 14 known NLRP3 interactors (16%) were discerned as high-confidence hits within the proximal proteomic data (**Figures 2D-H, Figure S4**) confirming the enrichment of NLRP3-associated proteins.

### Temporal tracking of the NLRP3 proximity proteome

To gain mechanistic and functional insights into the NLRP3 proximity proteome following nigericin stimulation at various intervals, we conducted a gene ontology-biological process (GO term-BP) enrichment analysis (**Figure 2I**). 13 shared terms, including viral translation, apoptotic cell death regulation, stress granule assembly and mitosis-associated processes, were significantly enriched across all four groups. In contrast to the unstimulated control, pathways related to “protein kinase D signalling” and “mRNA binding regulation” were uniquely present after nigericin-exposure, consistent with prior observations (Zhang et al., 2017). Intriguingly, the enrichment of “cellular response to arsenite ion” and “regulation of stress granule disassembly” terms suggest a potential functional association between inflammasome and stress granule formation. This connection was substantiated by previous reports of the stress granule protein DDX3X as a pivotal mediator, modulating between apoptotic ASC speck formation and pro-survival stress signalling (Fox and Man, 2019; Samir et al., 2019). In terms of GO term-molecular function (GO term-MF) enrichment analysis, “GTP-dependent protein binding” and “1- phosphatidylinostitol binding” were significantly enriched solely in the nigericin-60 min and nigericin- 90 min groups, suggesting that nigericin treatment might induce the recruitment of proteins associated with the above two biological functions (**Figures S5A-C**). GO term-cellular components (CC) analysis identified 16 common subcellular compartments or complexes, including COPII vesicle coat, ESCRT-0 complex, and chaperone complex. Notably, three endocytic recycling-associated complexes, including the endosome-associated recycling protein (EARP) complex, endosome to plasma transport vesicle and Golgi-associated retrograde protein (GARP) complex, were significantly enriched solely in nigericin-60 min and nigericn-90 min groups, suggesting that the NLRP3 complex may translocate to trans-Golgi network (TGN) or endosome upon stimulation (**Figure S5D**). Similarly, the compartments enrichment analysis identified another four complexes, including Dcp1-Dcp2, signal recognition receptor, MAD1 and the cohesin I complex, which were only captured and enriched in the last two timepoints of nigericin treatment (**Figures S6A-E**). We also performed tissue enrichment analysis, and not surprisingly, poor tissue specificity was observed in either unstimulated or stimulated groups (**Figure S5E**). To further explore the physiological and pathological biological processes, a reactome pathway analysis was performed on the proximity proteome. The top 20 significantly enriched pathways are shown as in **Figure S6F**. Notably, several innate immunity and virus infection associated pathways, such as interleukin-6 signalling, MHC class I complex, and PP2A-mediated dephosphorylation of key metabolic factors, were identified and enriched. Selected gene sets involved in the above pathways are plotted in **Figures S6G-H**.

### Localization analysis reveals translocation of NLRP3 upon stimulation

As the subcellular relocalization of NLRP3 and its underlying mechanism remain controversial (Hamilton and Anand, 2019), we performed the localization analysis to assess the spatial properties of NLRP3 inflammasome during complex assembly. After the filtering, normalization, and imputation, we compared the relative abundance of biotinylated proteins in the unstimulated (DMSO) and stimulated states (nigericin-90 min). The resulting volcano plot (**Figure S3F**) shows 103 proteins enriched in the right quadrant for the stimulated state, while 104 proteins enriched in the left quadrant for the resting state, with relatively stringent cut-off for fold change and false discovery rate (log2[FC]≥0.5 or ≤-0.5, FDR≤ 0.05). GO term-CC enrichment analysis was performed on the differentially enriched proteins in either condition. The proximity proteome under stimulated condition showed high spatial specificity for Golgi apparatus, with 24 proteins falling into the GO-CC term category-Golgi membranes (**Figures S3D-F, Table S2**). This is consistent with the previous observation that NLRP3 is activated and recruited to dispersed trans-Golgi networks (dTGN) in response to diverse stimuli (Bordon, 2019; Chen and Chen, 2018).

### Mapping stimulation-dependent NLRP3 interaction alterations using APEX2-PL and AP-MS

Modulation of the NLRP3 local milieu during the activation process was further examined using an integrated approach that combined NLRP3 APEX2 proximity labelling (PL) and affinity purification mass spectrometry (AP-MS) (**Figure 3A**). For APEX2 PL-MS, the SAINTexpress analysis identified 848 (DMSO) and 833 (Nig-90min) proximal proteins, while 484 (DMSO) and 366 potential interacting proteins were captured using the AP-MS method (**Figures 3A, B, Figures S7A-H**). There was relatively small number of overlapping proteins between APEX2 and AP-MS possibly as a result of the complementary nature of these two techniques (**Figures 3C, D**) and consistent with previous studies comparing BioID and AP-MS (Lambert et al., 2015). Additionally, we sought to compare the proximal proteome of NLRP3 under both resting and activation states. Following a rigorous filtration, seven known NLRP3-interacting proteins were differentially enriched. Of these, five proteins (NEK7, MAVS, PRKD1, PRKD2, and PRKD3, highlighted in red) enriched in the right quadrant are established positive regulators of NLRP3. In contrast, EIF2AK2 and LYN (highlighted in green) in the left quadrant are previously characterized inhibitory mediators. Interestingly, we noticed an increased labelling and capture of NLRP3 under stimulated conditions, likely due to oligomerization (**Figure 3E**).

**Figure 3.**
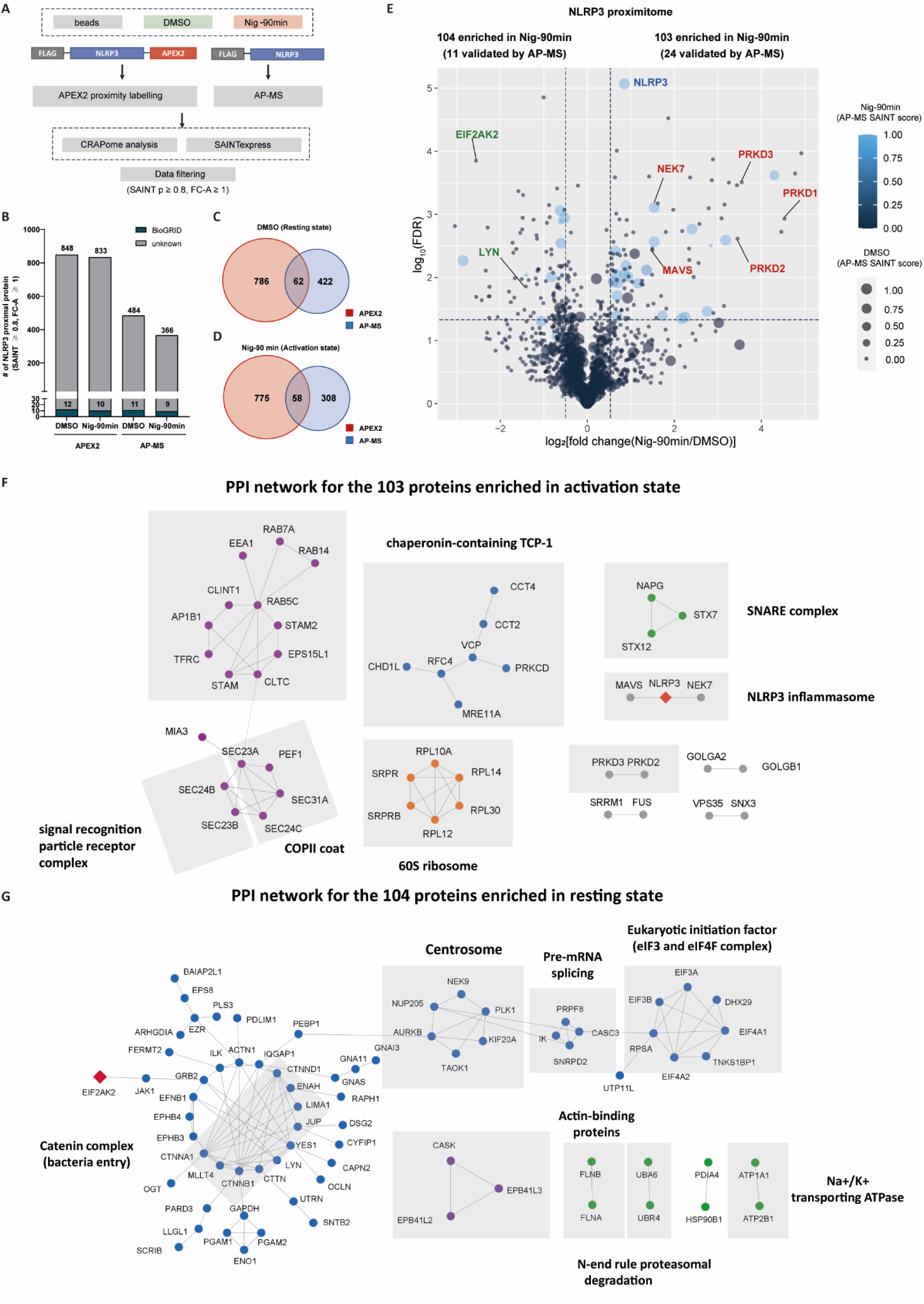
Mapping stimulation-dependent NLRP3 PPI alterations using APEX2-PL and AP-MS. (A) Schematic of the integrated approach combining APEX2 proximity labelling and AP-MS. (B) The number of known (BioGRID, blue) and newly discovered NLRP3-interacting proteins (grey) from APEX2 proximity labelling or AP-MS. (C-D) Venn diagrams showing the number of NLRP3 proximal proteins identified in APEX2 proximity labelling or AP-MS. (E) Volcano plot depicting the fold change for relative protein quantification of NLRP3-APEX2 proximity labelling samples between unstimulated (DMSO) and stimulated conditions (nigericin-90min). Data are shown as means of three independent samples. Non-adjusted unpaired two-tailed t-test was applied to the statistical analysis. Sizes denote the SAINT probability score for AP-MS during stimulation (nigericin-90min), and colours reflect the score in its unstimulated condition. (F-G) Protein-protein interaction network for NLRP3-APEX2 proximity labelled proteins enriched after stimulation with nigericin (F) or unstimulated as control (G). MCL algorithm was performed with the PPI scores from STRING database (confidence score threshold = 0.7). Grey lines denote the high-confidence protein-protein interaction curated from Reactome, CORUM and IntAct database. Cluster labels were added manually. n = 3 biological replicates.

To gain a visual understanding of the interaction networks within the NLRP3 proximity proteome and infer related biological insights, we performed a PPI network analysis. As for the nigericin stimulated condition (**Figure 3F)**, we identified protein complexes including the NLRP3 inflammasome, NEK7, MAVS, the SNARE complex, COPII vesicle coat, the signal recognition particle receptor complex and the protein kinase D signalling complex. **Figure 3G** highlights the interaction networks linked to the unstimulated state, in which the centrosome, the catenin complex, translational initiation factor, calcium dependent serine kinase and the N-end rule proteasomal pathway were significantly enriched.

### NLRP3-associated DUBs identified by proximity proteome-guided RNAi screening

The proximity proteome enabled us to capture several ubiquitin system components including 18 DUBs in total (USP9X, USP8, USP5, USP47, USP32, USP24, USP19, USP15, USP14, USP10, USP1, UCH-L1, PAN2, OTULIN, OTUID4, OTUB1, EIF3H and CYLD), which passed the stringent filtering criteria (SAINT probability > 0.8, FC-A ≥ 1). Among the 18 enriched DUBs, 9 were consistently present in all the four groups, and no marked difference was noted between the stimulated and unstimulated states (**Figures 4A-E**). To complement this, we carried out a DUB library RNAi screen in PMA-differentiated THP-1 cells, aiming to assess their influences on NLRP3 inflammasome priming and activation, reflected by IL-1β production. In total, knockdown of 25 DUBs (outliers that surpass mean ± 1 standard deviation, z score ≥1 or ≤-1, **Table S3**) were found to alter IL-1β levels in THP-1 cells treated with LPS or LPS /nigericin (**Figures 4F-G**). A combination of proximity proteomics and RNAi screening revealed 7 DUBs showing effects in both assays (USP9X, USP8, USP14, USP10, UCH-L1, OTUB1 and OTUD4) (**Figure 4H**). USP9X, USP14, and OTUB1 have already been reported to modulate the NLRP3 inflammasome (Hai et al., 2022; Xiang et al., 2023), suggesting additional NLRP3-associated DUBs (USP8, USP10, OTUD4 and UCH-L1).

**Figure 4.**
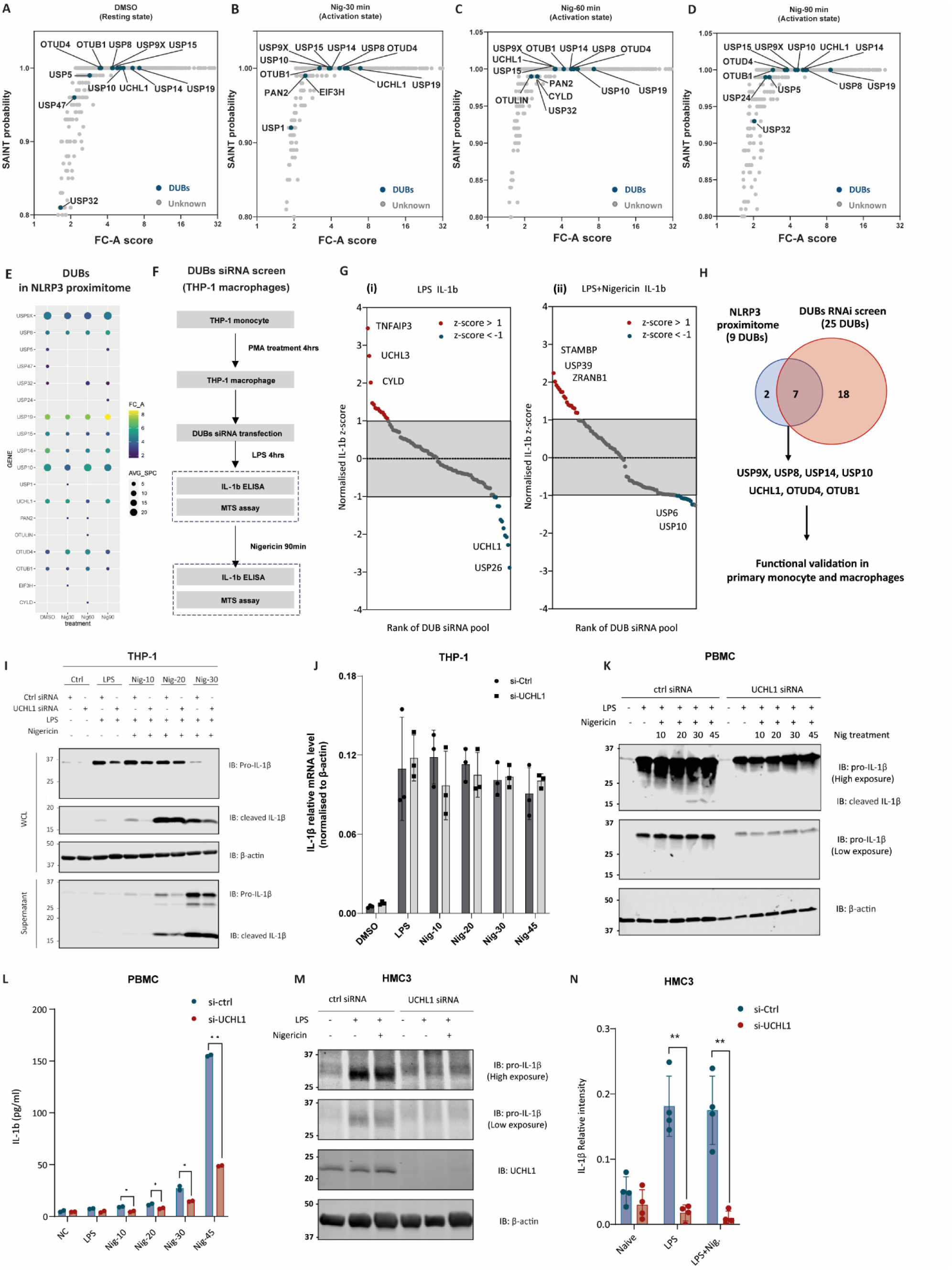
Proximity proteome-guided RNAi screen identifies deubiquitinases that are involved in IL-1β production. (A-D) APEX2 proximity labelling identifies DUBs in the local milieu of NLRP3 upon nigericin stimulation. SAINT probability scoring of the proteomic data for NLRP3-APEX2 compared to the negative beads control. Proteins passing the cut off (SAINT probability score ≥ 0.8 and FC-A score ≥ 1) are considered as high confidence proximal proteins for NLRP3. Deubiquitinases are coloured in blue and all other proteins are shown in grey. (E) Dot plots showing the DUBs in the NLRP3 proximity proteome upon nigericin stimulation for different time points as indicated. Sizes denote the average spectral counts and colours denote the FC-A score. (F) Schematic of experimental design for the DUB siRNA library screen. (G) z-scores distribution of the IL-1β in the supernatant for individual siRNAs. The RNAi screen is carried out in PMA-differentiated THP-1 cells upon LPS (i) or LPS+NIG treatment (ii), using ELISA for the measurement of IL-1β in the supernatant. Hits with a z-score of ≤ -1 or ≥1 are chosen as candidates for further validation. (H) Venn diagrams showing the intersection between the hits identified through APEX2-PL and siRNA screen. (I-J) Immunoblot and RT-qPCR analysis of whole cell lysates and supernatants from THP-1 cells transfected with non-targeting control siRNA, or UCH-L1-specific siRNA. Cells were differentiated with PMA and primed with LPS (1 μg/mL) for 4 hours, followed by the treatment of nigericin (10 μM) for different timepoints as indicated. (K-L) PBMCs were transfected with non-targeting control, or UCH-L1-specific siRNA, stimulated with LPS for 4 hours and nigericin for different timepoints as indicated. (K) Immunoblot of cell lysates from PBMCs. (L) Production of IL-1β in the supernatants from PBMCs as measured by ELISA. *p < 0.05, **p < 0.01 (unpaired two-sided t-test). The results are representative of 3 independent experiments. (M- N) Immunoblot of cell lysates from HMC3 cells transfected with non-targeting control siRNA or UCH- L1-specific siRNA. Cells were primed with LPS (1 μg/mL) for 4 hours, followed by the treatment of nigericin (10 μM) for 45 minutes. (N) IL-1β bands were quantified using GelAnalyzer and normalised using β -actin bands. Error bars show the standard deviation of the mean. **p < 0.01 (unpaired two- sided t-test).

### UCH-L1 interacts with the NACHT domain of NLRP3 and affects IL-1β production

Amongst these DUBs, UCH-L1 consistently affected IL-1β levels and showed high NLRP3 proximity scores. Notably, the knockdown of UCH-L1 did not alter pro-IL-1β mRNA expression, while it decreased protein levels and production of the mature form in THP-1 monocytes (**Figures 4I, J**) and human PBMCs (**Figures 4K, L**). Expression of IL-6 and TNFα were also slightly suppressed in the absence of UCH-L1 (**Figure S8B, C**), indicating that UCH-L1 might also play a role in the NF-κB-regulated priming stage. As UCH-L1 is highly abundant in cells associated with the central nervous system, we sought to investigate its role in brain resident immune cells, such as microglia, which also express NLRP3 inflammasome components (**Figure S9**). Similarly to THP-1 and PBMC cells, UCH- L1 deficiency markedly suppressed IL-1β production in the human microglial cell line HMC3 (**Figures 4M, N**).

The NLRP3-UCH-L1 interaction was identified through NLRP3-APEX2 proximity labelling (**Figure 5A**) and confirmed by immunoprecipitation (**Figure 5B**) in HEK293 cells. Notably, the interaction between NLRP3 and UCH-L1 decreases upon nigericin stimulation (**Figure 5B**). To further determine which domain within NLRP3 interacts with UCH-L1, we overexpressed flag-tagged full-length and truncated NLRP3 in HEK293T cells. UCH-L1 interacted with the full-length NLRP3 and NACHT domain of NLRP3 (**Figure 5C**). These results imply that the NACHT domain of NLRP3 directly or indirectly provides a binding site for UCH-L1. Additionally, immunofluorescence revealed the colocalization of NLRP3 with UCH-L1 in hiPSC-derived microglial cells (**Figures 5D, E**).

**Figure 5.**
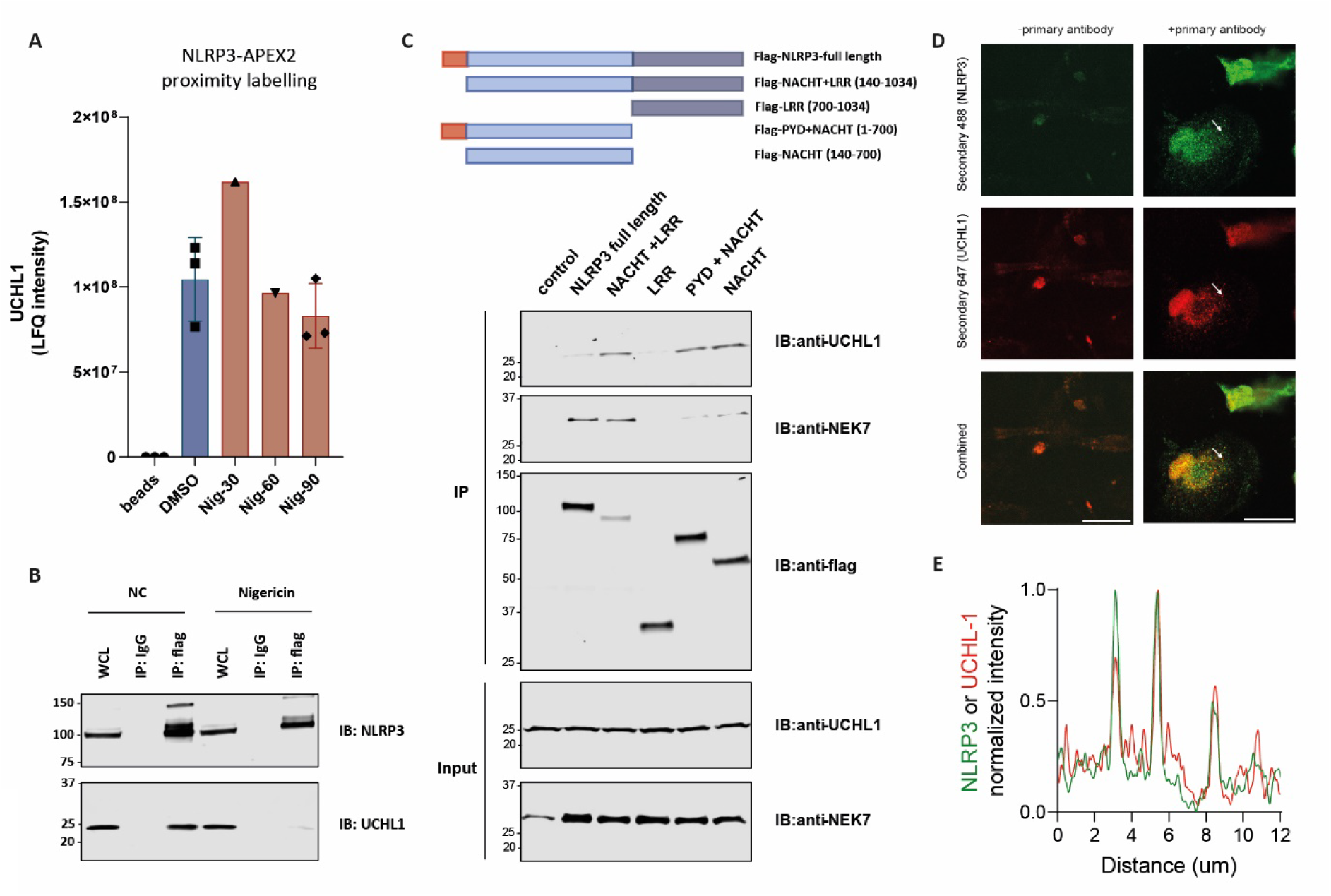
UCH-L1 interacts with the NACHT domain of NLRP3. (A) The raw LFQ Intensity of UCH-L1 protein identified in the APEX2 proximity labelling performed in this study across all the samples including beads Control, DMSO Control and Nigericin treatment at different time points. Experiment was performed in triplicate except of Nig-30 and Nig-60 where only 1 replicant was performed. (B) Flp-In T-Rex 293 NLRP3-flag stable cells were induced with tetracycline overnight. Afterwards, cells were treated with DMSO or Nigericin for 90 minutes. NLRP3- flag was pulled down using anti-flag antibody or IgG and immunoblotted with indicated antibodies. (C) Schematic representation of the N-terminal Flag-tagged NLRP3 domain constructs: full length NLRP3, NACHT and LRR domain NLRP3 (Flag-NACHT+LRR, amino acids 140-1034), LRR domain of NLRP3 (Flag-LRR, amino acids 700-1034), pyrin and NACHT domain of NLRP3 (Flag- PYD+NACHT, amino acids 1-700), NACHT domain of NLRP3 (Flag-NACHT, amino acids 140-700). HEK293T cells were transfected with the appropriate NLRP3 plasmid using Lipofectamine 3000. Cells were cultured for 24 hours before they were lysed. NLRP3 domain-flag was pulled down using anti- flag antibody and immunoblotted with indicated antibodies. (D) Fluorescence confocal images of hiPSC-derived microglial cells stained for UCHL-1 and NLPR3. UCHL-1 and NLRP3 show colocalization, but only when cells were stained with primary antibodies for UCHL-1 and NLRP3 (right column). No colocalization is shown when cells are not stained with primary antibodies against UCHL- 1 and NLRP3 (left column). (E) Line profile showing the min-max normalized intensity of UCHL-1 (red) and NRLP3 (green) along and in the direction of the white arrow indicated in (D). The peaks in normalized intensity indicate colocalization of NLRP3 and UCHL-1 in enriched areas (or ‘specks’) of NLRP3.

### UCH-L1 catalytic inhibition abrogates ASC assembly, NLRP3 inflammasome activation and IL- 1β processing

To investigate how UCH-L1 modulates IL-1β processing and release, we selected a potent inhibitor- IMP-1711-S (small molecule 27 in patent (Mark Kemp, 2017), **Figure 6A**) (Panyain et al., 2020; Panyain et al., 2021) and tested its effect on NLRP3 inflammasome activation in THP-1, mouse bone marrow-derived macrophages (BMDMs), human PBMCs, and hiPSC-derived microglial cells. Cells were primed with LPS, pre-treated with IMP-1711-S and stimulated with the NLRP3 activator nigericin. We observed that the treatment with IMP-1711-S inhibited the production of IL-1β in a concentration-dependent manner in PMA-differentiated THP-1 (**Figure 6B**) and hiPSC-derived microglia (**Figures 6K, L**). Both, caspase-1 p10 and cleaved IL-1β levels were reduced in supernatants from IMP-1711-S-treated THP-1 cells (**Figures 6C-E**), suggesting that the inhibitor exerts its effect on the activation of the caspase-1 and IL-1β processing stage. IMP-1711-S potently inhibited the release of lactate dehydrogenase (LDH) and blocked pyroptotic cell death induced by nigericin (**Figures 6F, G**). Additionally, IMP-1711-S treatment did not affect pro-caspase-1 or pro-IL-1β expression in lysates in the priming stage with LPS stimulation (**Figure 6C**). Further investigation of ASC oligomerization in THP-1 cells through crosslinking of the insoluble lysate fraction revealed that while IMP-1711-S had no impact on ASC expression in cell lysates, it substantially reduced oligomerized ASC levels post LPS and nigericin stimulation (**Figure 6H**). Additionally, in HEK293 cells, IMP-1711-S markedly downregulated the formation of NLRP3 specks in a concentration-dependent manner (**Figures 6I, J**). Our findings underscore that UCH-L1 inhibition impedes NLRP3 inflammasome activation by disrupting NLRP3 assembly and the speck formation, ultimately leading to decreased IL-1β production (**Figure 7**).

**Figure 6.**
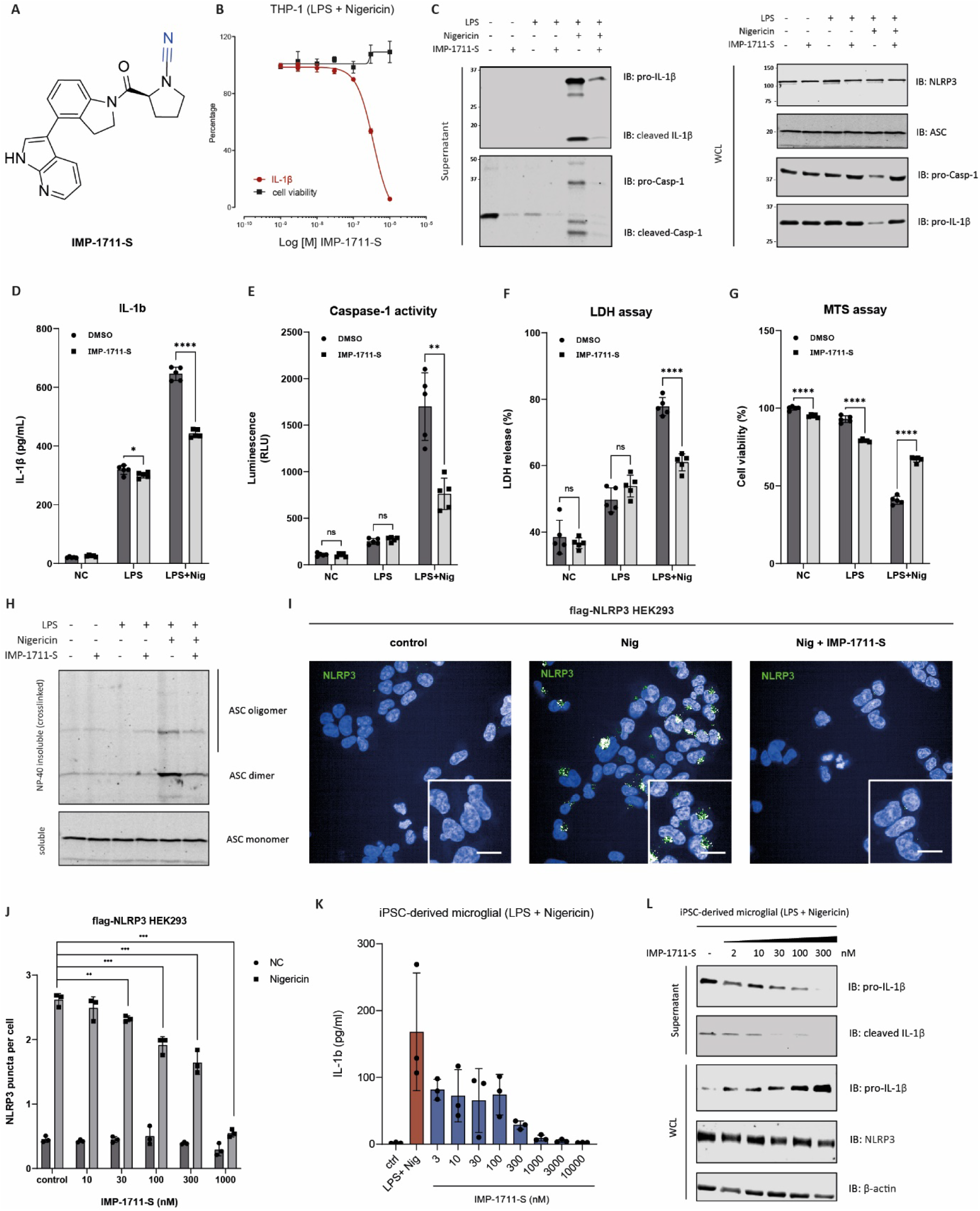
UCH-L1 inhibitor blocks ASC/NLRP3 assembly and IL-1β production. (A) UCH-L1 inhibitor IMP-1711-S structure. (B) Red, production of IL-1β from PMA-differentiated THP-1 cells stimulated with LPS and nigericin and treated with IMP-1711-S (3-10,000 nM) as measure by ELISA (red). Black, cell viability measured by MTS assay. Data are expressed as mean ± s.d., n = 3 biologically independent samples. (C-G) THP-1 cells were differentiated with PMA and primed with LPS (1 μg/mL) for 4 hours, followed by the treatment of nigericin (10 μM) for 45 minutes. DMSO or IMP-1711-S (0.3 μM) was added at the same time with LPS. (C) Immunoblot analysis of whole cell lysates and supernatants from THP-1 cells treated with DMSO or IMP-1711-S. (D-E) Supernatants were collected for the measurement of IL-1β (D), Caspase-1 activity (E) from 3 independent experiments. *p < 0.05, **p < 0.01, ***p < 0.001, ****p < 0.0001 (unpaired two-sided t-test). Additionally, the cell viability was measured using LDH assay (F) and MTS assay (G). (H) Immunoblot analysis of ASC polymerization upon treatment of DMSO or IMP-1711-S. (I-J) Immunofluorescence imaging of flag-NLRP3 HEK293 stable cells treated with DMSO or IMP-1711-S (300nM), followed by the stimulation with nigericin. Scale bars, 10 μm. (J) Quantitative analysis of the NLRP3 puncta formation upon nigericin treatment in the presence of DMSO or different doses of IMP-1711-S (0.001- 1μM) as indicated. **p < 0.01, ***p < 0.001. (K) IL-1β levels were determined using ELISA from the supernatant of fully differentiated hiPSC-derived microglia. Cells were primed with LPS (1 μg/mL) for 4 hours, followed by the treatment of nigericin (10 μM) for 45 minutes. IMP-1711-S inhibitor was added simultaneously with LPS at different concentrations. (L) Immunoblot analysis of whole cell lysates and supernatants from hiPSC-derived microglia treated with DMSO or different concentrations of IMP-1711-S as indicated.

**Figure 7.**
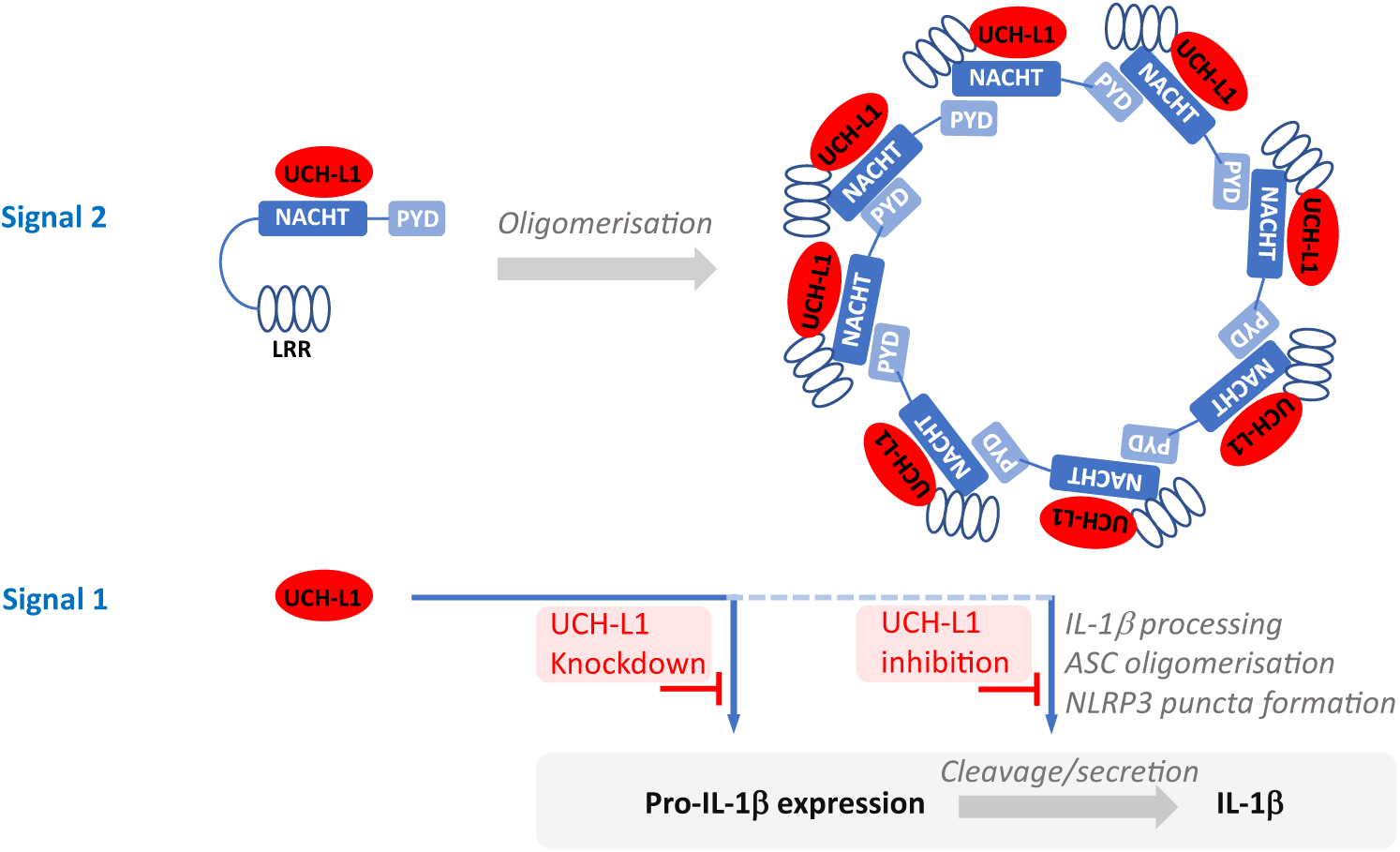
Model illustrating multi-functional roles of UCH-L1 in the NLRP3-mediated IL-1β expression, processing and production. UCH-L1 was shown previously to play a role in NF-κB signalling, affecting pro-IL-1β expression (Zhang et al., 2021) in the context of signal 1. UCH-L1 deletion or pharmacological inhibition interferes with IL-1β production through processes of signal 2 (nigericin) involving ASC oligomerisation, thereby blocking IL-1β processing and secretion.

## DISCUSSION

The NLRP3 inflammasome is central to innate immunity, and a detailed understanding of its molecular dynamics and interactome is highly sought after. We found that NLRP3 tolerates C-terminally tagged APEX2, retaining its competence to biotinylate and capture known interacting partners of endogenous NLRP3. Using complementary techniques, such as APEX2 PL-MS, AP-MS, and siRNA screening, we traced the time-resolved PPI networks local to the NLRP3 inflammasome upon stimulation. This approach offers a resource-rich map for further investigations of the molecular intricacies of NLRP3 inflammasome activation and revealed that UCH-L1 is involved in regulating IL-1β production.

### Deconvolution of the NLRP3-APEX2 proximity proteome

A critical issue related to proximity labelling experiments is discrimination between true interactors and background noise. Proteins captured by NLRP3-APEX2 proximity labelling can be categorized into four groups: Type I-True interacting proteins that physically interact with NLRP3 and or other components of the complex; Type II-Compartmental or proximal proteins in the local milieu of NLRP3 reflecting transient interactions; Type III-non-specific proteins that stick to streptavidin beads; Type IV-physically non-existing proteins detected in error. Type I and type II provide insights into the PPI network and local milieu of NLRP3, but type III and type IV are considered false positives. To establish a high confidence proximity proteome resource for NLRP3, a variety of approaches were used to filter out type III and type IV contaminant proteins. A Bayesian model-based algorithm (SAINTexpress) was applied to analyze the proximity labelling data with a SAINT probability score threshold of 0.8 and fold change ≥ 2, yielding 800-1100 proximal proteins per group (**Figures 2C, E-H**). Furthermore, the unstimulated state was used as a control, which helped us distinguishing the differentially regulated interacting partners upon stimulation with nigericin, still yielding more than 800 proteins per biological condition, consistent with the APEX2 enzyme affording a biotin labelling radius of ∼ 10 nm in live cells (Qin et al., 2021). With a stringent cut-off, our data identified 103 high-confidence putative NLRP3-associated proteins enriched under activated conditions and 104 proteins enriched under inactivated conditions. These included known interactors, such as NEK7, MAVS, PRKD1/2/3, LYN and EIF2AK2 (**Figure 3E**). We note that spatial references have been shown to enhance the deconvolution of PL data. However, in our study, this was not applied as NLRP3 resides in multiple compartments and translocates to distinct organelles upon activation (Hamilton and Anand, 2019). By integrating PL data with AP-MS and siRNA screening, we established an alternative method for validating NLRP3-associated proteins.

### UCH-L1 is an essential mediator of NLRP3/ASC assembly and IL-1b production

Ubiquitin modification controls multiple elements of the NLPR3 activation process and IL-1β production (Lopez-Castejon, 2020). Through a DUB siRNA screen, we pinpointed four deubiquitylating enzymes, TNFAIP3, CYLD, USP7, and STAMBP, already been linked to IL-1β signalling pathways (**Figure 4G)** (Bednash et al., 2021; Duong et al., 2015; Palazon-Riquelme et al., 2018; Vande Walle et al., 2014; Yang et al., 2020). However, the intricate nature of IL-1β processing, spanning multiple regulatory stages, poses a challenge to precisely attribute the functional impacts of each DUB. For instance, UCH-L3 and L5 were suggested to affect inflammasome activation and IL-1β production, although in the context of infection (Kummari et al., 2015; Qu et al., 2020; Ramachandran et al., 2021). UCH-L3 may act in the priming stage, whereas UCH-L5’s role maybe less clear outside the context of pathogen-driven NLRP3 inflammasome activation, consistent with the former, but not the latter, being detected in our screen (**Fig. 4G**).

Here, we identified UCH-L1 as an NLRP3 interacting modulator of the inflammasome. It is a member of the ubiquitin C-terminal hydrolase (UCH) family involved in neurodegeneration and cancer metastasis (Ham et al., 2021; Liu et al., 2002; Mondal et al., 2022b). UCH-L1 is highly expressed in the central nervous system, with low expression levels in most other tissues under normal physiology. Dysregulation of UCH-L1 expression and mutations have been implicated in multiple neurodegenerative diseases, such as Parkinson’s disease, Huntington’s disease, and recessive hereditary spastic paraplegia (SPG79) (Mondal et al., 2022a). Also, several studies linked UCH-L1 to inflammation or innate immunity. For instance, UCH-L1 deficiency markedly reduces the secretion of pro-inflammatory cytokines, notably IL-6 and TNFα (Zhang et al., 2021). LDN57444, an isatin O-acyl oxime small molecule inhibitor, has demonstrated efficacy in ameliorating LPS-induced sepsis in *in vivo* models (Zhang et al., 2021), although other studies have cast doubt on the specificity or efficacy of this compound against UCH-L1 (Panyain et al., 2020). Another study reported that UCH-L1 promoted M1 macrophages polarization via the PI3K/Akt signalling pathway (Huang et al., 2022). Consistent with previous findings, UCH-L1 knockdown or inhibition yielded a significant decrease in pro-inflammatory cytokine expression and secretion including IL-1β, IL-6 and TNFα in both, THP-1 and human primary monocytes, reflecting interference in upstream signalling affecting the priming stage (**Figures 4I-L, Figures S8, S10**). Furthermore, mRNA levels of IL-1β remain unchanged in the absence of UCH-L1, suggesting that UCH-L1 affects IL-1β at the post-transcriptional or protein level (**Figure 4J**). Notably, pharmacological inhibition of UCH-L1 affected ASC/NLRP3 assembly and IL-1β production, which are part of the second inflammasome induction signal (**Figure 6**). Although the small molecule inhibitor IMP-1711-S altered secreted active caspase-1 levels (**Figure 6E**), it does not seem to cross-react or inhibit caspases directly (Panyain et al., 2020). Caspase-1 inhibitors also show a different specificity profile towards TNFα release (Cao et al., 2022). UCH-L1 inhibition interferes with upstream events, such as NLRP3 puncta formation and ASC oligomerization (**Figure 6**), the latter of which is phenocopied by UCH-L1 knockdown (**Figure S11**). UCH-L1, found to be bound to the NACHT domain of NLRP3, may act as a gating/editing factor to control the ubiquitylation status of multiple components, such as NLRP3, ASC, Caspase-1 and IL-1β, required for NLRP3-ASC inflammasome complex assembly (**Figure 7**). Preliminary results indicate that NLRP3’s half-live is dependent on the ubiquitin-proteasome pathway (**Figure S12**). However, NLRP3 associated ubiquitylation appears to be minimally altered by UCH-L1, and NLRP3 protein levels are unaffected (**Figures S12, 13**), suggesting a more structural role for UCH-L1 and/or its modulation of NLRP3 ubiquitination during assembly rather than controlling turnover. Though speculative at this stage, such a putative structural and ubiquitylation editing “gate keeping” process maybe in analogy to the role of UCH-L5/UCH37 in the 26S proteasome complex, required for substrate deubiquitylation and ubiquitin recycling prior to substrate unfolding by the 19S and entry into the 20S for destruction (Song et al., 2021; Vander Linden et al., 2015).

In conclusion, in this study, we provide a detailed NLRP3 inflammasome interaction map at different physiological stages. UCH-L1 was identified and validated as an essential regulator involved in NLRP3 inflammasome activation and IL-1β production, affected by UCH-L1 chemical inhibition and phenocopied by UCH-L1 deletion. Our results may have translational implications in exploring pharmacological inhibition of UCH-L1 in neuroinflammation-associated pathology, which could include dementia.

## STAR★Methods

### Key resources table

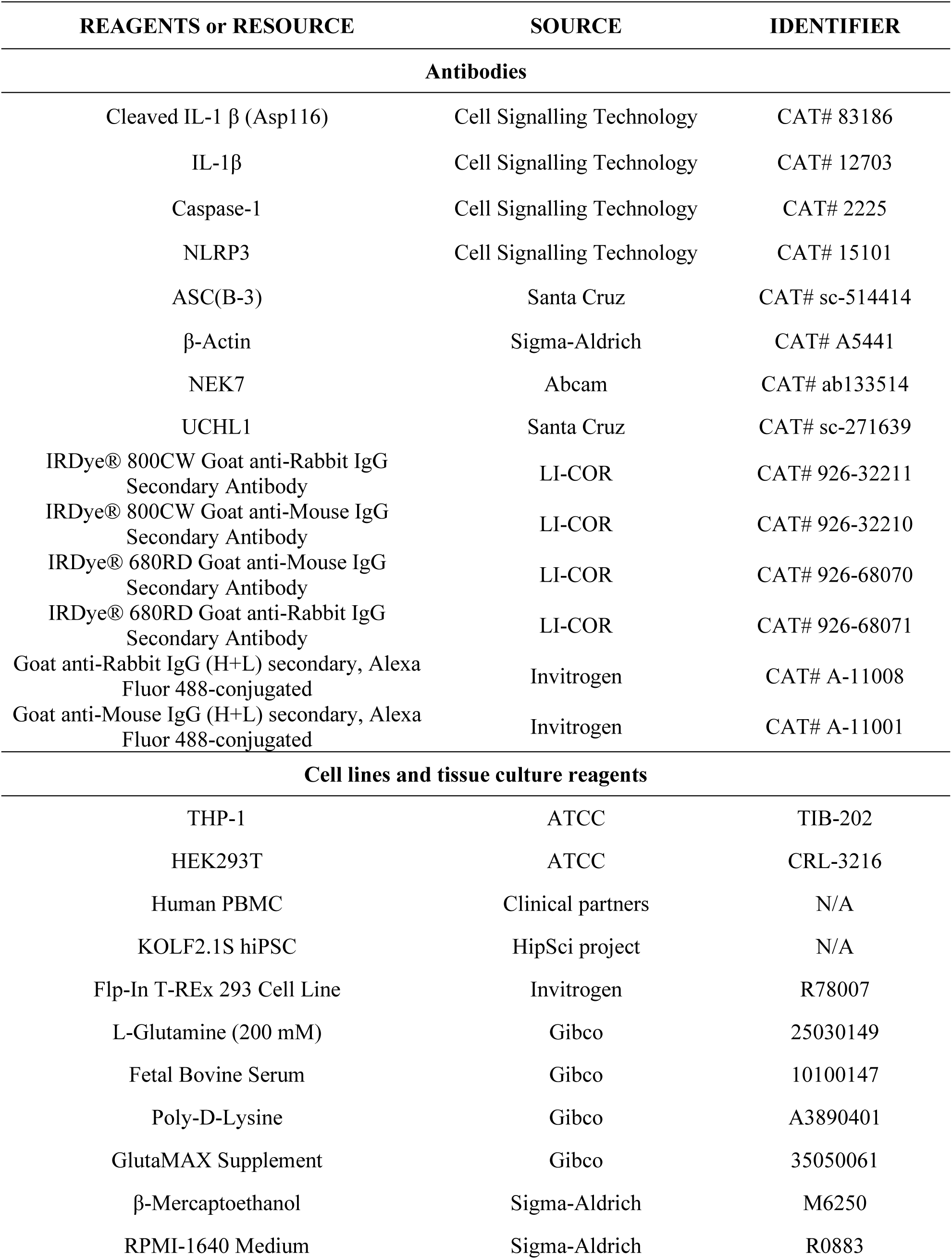
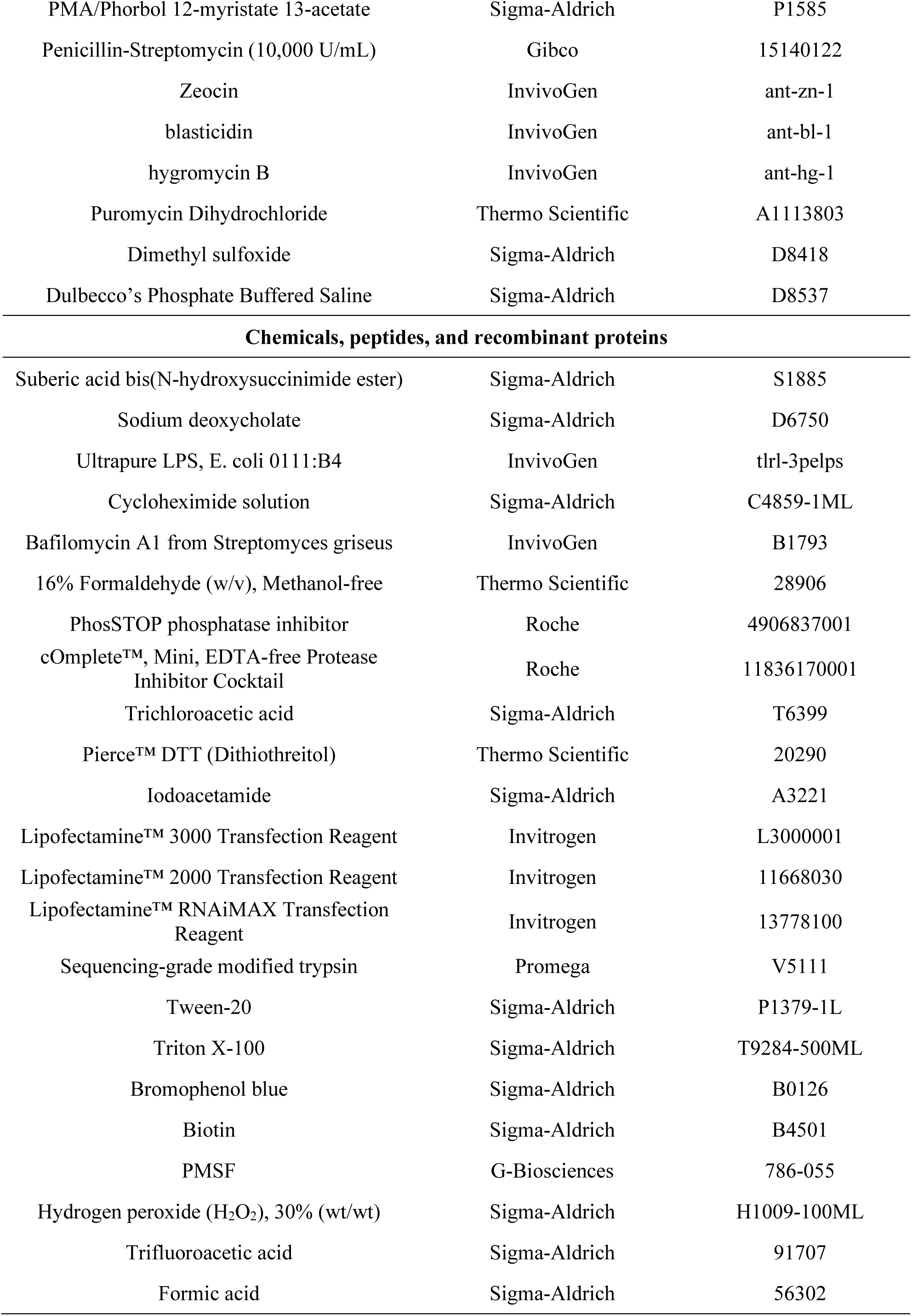
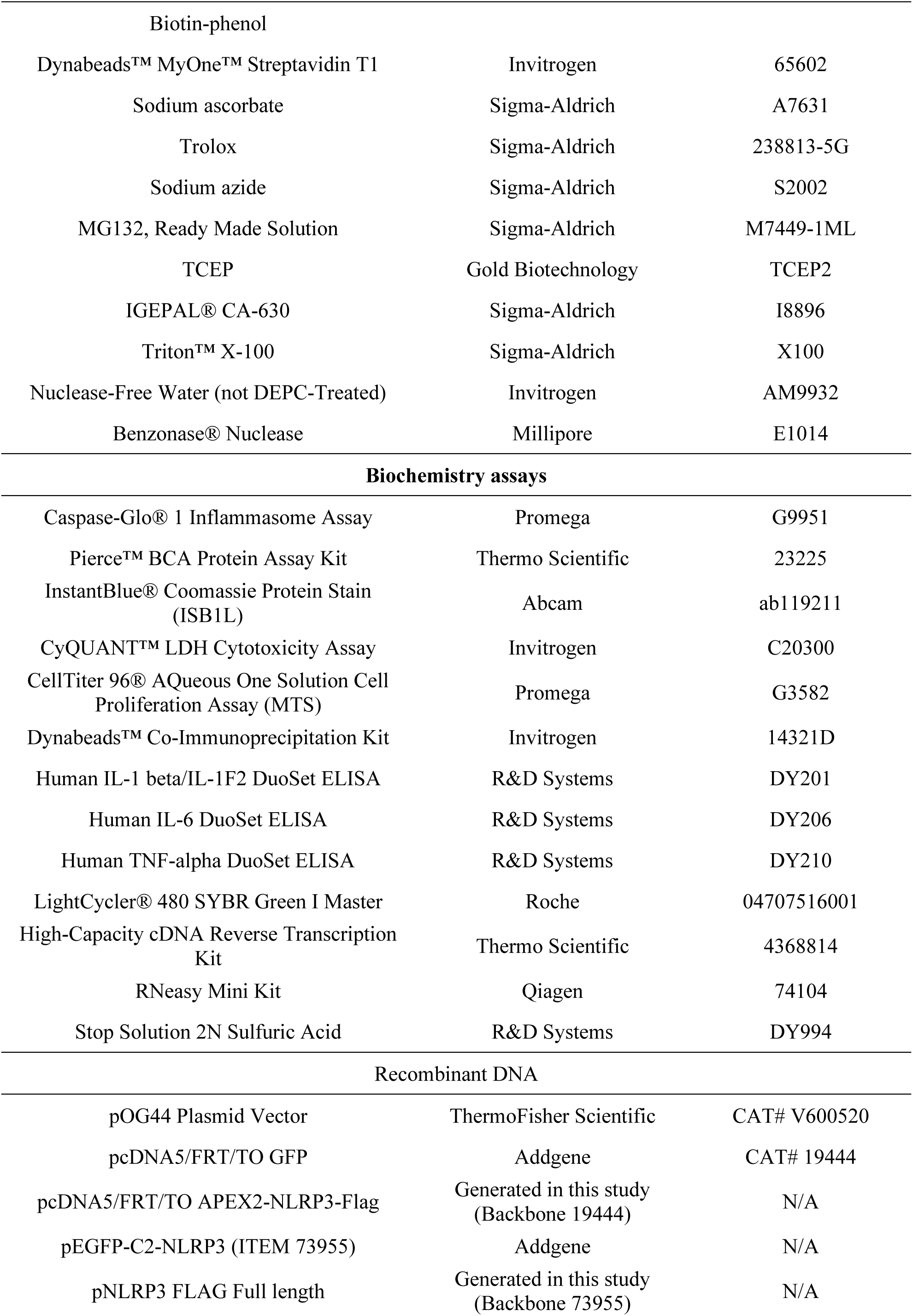
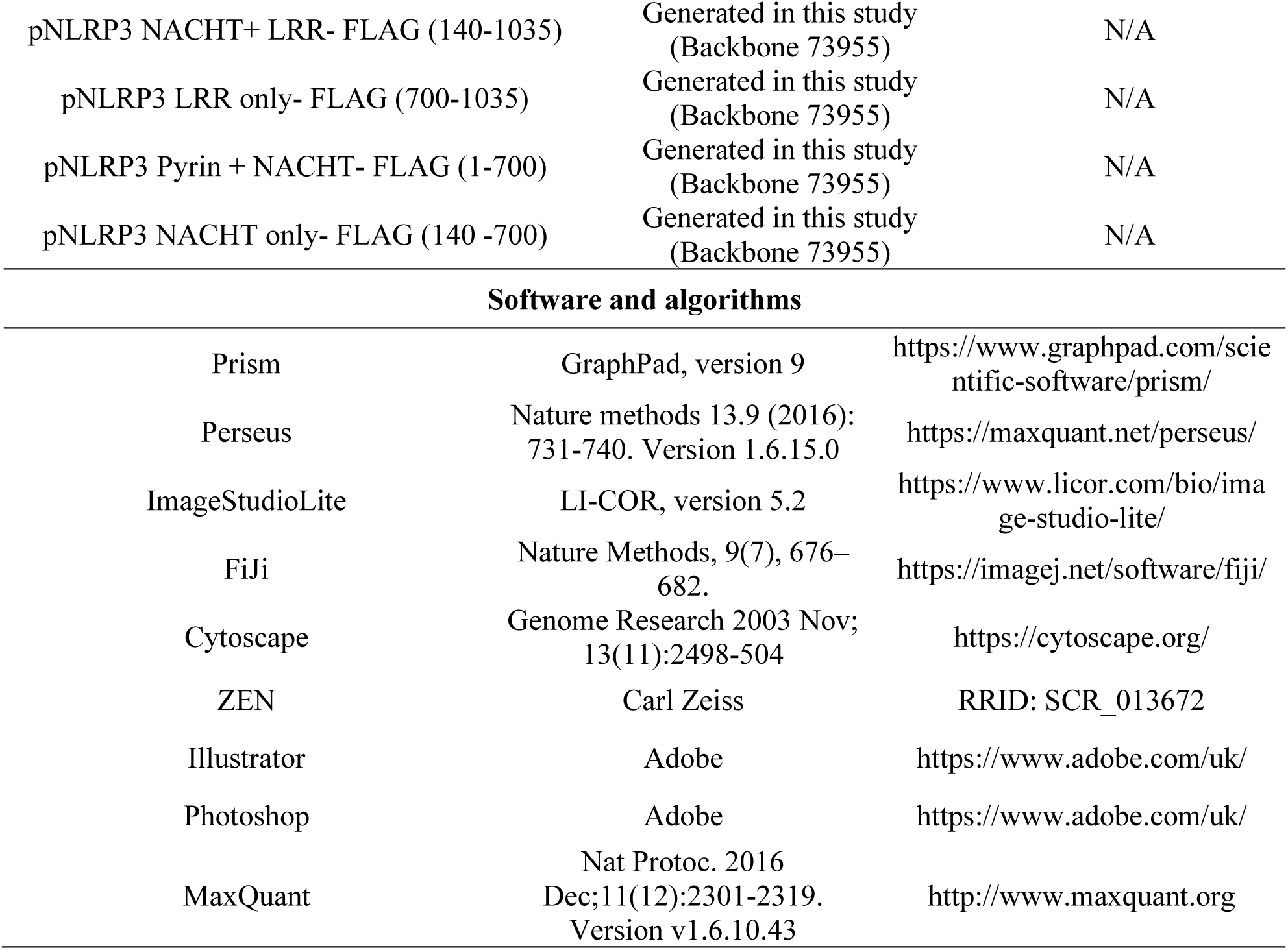

| REAGENTS or RESOURCE | SOURCE | IDENTIFIER |
| --- | --- | --- |
| **Antibodies** |  |  |
| Cleaved IL-1 $\beta$ (Asp116) | Cell Signalling Technology | CAT# 83186 |
| IL-1β | Cell Signalling Technology | CAT# 12703 |
| Caspase-1 | Cell Signalling Technology | CAT# 2225 |
| NLRP3 | Cell Signalling Technology | CAT# 15101 |
| ASC(B-3) | Santa Cruz | CAT# sc-514414 |
| β-Actin | Sigma-Aldrich | CAT# A5441 |
| NEK7 | Abcam | CAT# ab133514 |
| UCHL1 | Santa Cruz | CAT# sc-271639 |
| IRDye® 800CW Goat anti-Rabbit IgG Secondary Antibody | LI-COR | CAT# 926-32211 |
| IRDye® 800CW Goat anti-Mouse IgG Secondary Antibody | LI-COR | CAT# 926-32210 |
| IRDye® 680RD Goat anti-Mouse IgG Secondary Antibody | LI-COR | CAT# 926-68070 |
| IRDye® 680RD Goat anti-Rabbit IgG Secondary Antibody | LI-COR | CAT# 926-68071 |
| Goat anti-Rabbit IgG (H+L) secondary, Alexa Fluor 488-conjugated | Invitrogen | CAT# A-11008 |
| Goat anti-Mouse IgG (H+L) secondary, Alexa Fluor 488-conjugated | Invitrogen | CAT# A-11001 |
| **Cell lines and tissue culture reagents** |  |  |
| THP-1 | ATCC | TIB-202 |
| HEK293T | ATCC | CRL-3216 |
| Human PBMC | Clinical partners | N/A |
| KOLF2.1S hiPSC | HipSci project | N/A |
| Flp-In T-REx 293 Cell Line | Invitrogen | R78007 |
| L-Glutamine (200 mM) | Gibco | 25030149 |
| Fetal Bovine Serum | Gibco | 10100147 |
| Poly-D-Lysine | Gibco | A3890401 |
| GlutaMAX Supplement | Gibco | 35050061 |
| β-Mercaptoethanol | Sigma-Aldrich | M6250 |
| RPMI-1640 Medium | Sigma-Aldrich | R0883 |
| PMA/Phorbol 12-myristate 13-acetate | Sigma-Aldrich | P1585 |
| Penicillin-Streptomycin (10,000 U/mL) | Gibco | 15140122 |
| Zeocin | InvivoGen | ant-zn-1 |
| blasticidin | InvivoGen | ant-bl-1 |
| hygromycin B | InvivoGen | ant-hg-1 |
| Puromycin Dihydrochloride | Thermo Scientific | A1113803 |
| Dimethyl sulfoxide | Sigma-Aldrich | D8418 |
| Dulbecco's Phosphate Buffered Saline | Sigma-Aldrich | D8537 |
| **Chemicals, peptides, and recombinant proteins** |  |  |
| Suberic acid bis(N-hydroxysuccinimide ester) | Sigma-Aldrich | S1885 |
| Sodium deoxycholate | Sigma-Aldrich | D6750 |
| Ultrapure LPS, E. coli 0111:B4 | InvivoGen | tlrl-3pelps |
| Cycloheximide solution | Sigma-Aldrich | C4859-1ML |
| Bafilomycin A1 from Streptomyces griseus | InvivoGen | B1793 |
| 16% Formaldehyde (w/v), Methanol-free | Thermo Scientific | 28906 |
| PhosSTOP phosphatase inhibitor | Roche | 4906837001 |
| cOmplete™, Mini, EDTA-free Protease Inhibitor Cocktail | Roche | 11836170001 |
| Trichloroacetic acid | Sigma-Aldrich | T6399 |
| Pierce™ DTT (Dithiothreitol) | Thermo Scientific | 20290 |
| Iodoacetamide | Sigma-Aldrich | A3221 |
| Lipofectamine™ 3000 Transfection Reagent | Invitrogen | L3000001 |
| Lipofectamine™ 2000 Transfection Reagent | Invitrogen | 11668030 |
| Lipofectamine™ RNAiMAX Transfection Reagent | Invitrogen | 13778100 |
| Sequencing-grade modified trypsin | Promega | V5111 |
| Tween-20 | Sigma-Aldrich | P1379-1L |
| Triton X-100 | Sigma-Aldrich | T9284-500ML |
| Bromophenol blue | Sigma-Aldrich | B0126 |
| Biotin | Sigma-Aldrich | B4501 |
| PMSF | G-Biosciences | 786-055 |
| Hydrogen peroxide (H<sub>2</sub>O<sub>2</sub>), 30% (wt/wt) | Sigma-Aldrich | H1009-100ML |
| Trifluoroacetic acid | Sigma-Aldrich | 91707 |
| Formic acid | Sigma-Aldrich | 56302 |
| Biotin-phenol |  |  |
| Dynabeads™ MyOne™ Streptavidin T1 | Invitrogen | 65602 |
| Sodium ascorbate | Sigma-Aldrich | A7631 |
| Trolox | Sigma-Aldrich | 238813-5G |
| Sodium azide | Sigma-Aldrich | S2002 |
| MG132, Ready Made Solution | Sigma-Aldrich | M7449-1ML |
| TCEP | Gold Biotechnology | TCEP2 |
| IGEPAL® CA-630 | Sigma-Aldrich | I8896 |
| Triton™ X-100 | Sigma-Aldrich | X100 |
| Nuclease-Free Water (not DEPC-Treated) | Invitrogen | AM9932 |
| Benzonase® Nuclease | Millipore | E1014 |
| Biochemistry assays |  |  |
| Caspase-Glo® 1 Inflammasome Assay | Promega | G9951 |
| Pierce™ BCA Protein Assay Kit | Thermo Scientific | 23225 |
| InstantBlue® Coomassie Protein Stain (ISB1L) | Abcam | ab119211 |
| CyQUANT™ LDH Cytotoxicity Assay | Invitrogen | C20300 |
| CellTiter 96® AQueous One Solution Cell Proliferation Assay (MTS) | Promega | G3582 |
| Dynabeads™ Co-Immunoprecipitation Kit | Invitrogen | 14321D |
| Human IL-1 beta/IL-1F2 DuoSet ELISA | R&D Systems | DY201 |
| Human IL-6 DuoSet ELISA | R&D Systems | DY206 |
| Human TNF-alpha DuoSet ELISA | R&D Systems | DY210 |
| LightCycler® 480 SYBR Green I Master | Roche | 04707516001 |
| High-Capacity cDNA Reverse Transcription Kit | Thermo Scientific | 4368814 |
| RNeasy Mini Kit | Qiagen | 74104 |
| Stop Solution 2N Sulfuric Acid | R&D Systems | DY994 |
| Recombinant DNA |  |  |
| pOG44 Plasmid Vector | ThermoFisher Scientific | CAT# V600520 |
| pcDNA5/FRT/TO GFP | Addgene | CAT# 19444 |
| pcDNA5/FRT/TO APEX2-NLRP3-Flag | Generated in this study (Backbone 19444) | N/A |
| pEGFP-C2-NLRP3 (ITEM 73955) | Addgene | N/A |
| pNLRP3 FLAG Full length | Generated in this study (Backbone 73955) | N/A |
| pNLRP3 NACHT+ LRR- FLAG (140-1035) | Generated in this study<br>(Backbone 73955) | N/A |
| pNLRP3 LRR only- FLAG (700-1035) | Generated in this study<br>(Backbone 73955) | N/A |
| pNLRP3 Pyrin + NACHT- FLAG (1-700) | Generated in this study<br>(Backbone 73955) | N/A |
| pNLRP3 NACHT only- FLAG (140 -700) | Generated in this study<br>(Backbone 73955) | N/A |
| Software and algorithms |  |  |
| Prism | GraphPad, version 9 | https://www.graphpad.com/scientific-software/prism/ |
| Perseus | Nature methods 13.9 (2016): 731-740. Version 1.6.15.0 | https://maxquant.net/perseus/ |
| ImageStudioLite | LI-COR, version 5.2 | https://www.licor.com/bio/image-studio-lite/ |
| FiJi | Nature Methods, 9(7), 676–682. | https://imagej.net/software/fiji/ |
| Cytoscape | Genome Research 2003 Nov; 13(11):2498-504 | https://cytoscape.org/ |
| ZEN | Carl Zeiss | RRID: SCR_013672 |
| Illustrator | Adobe | https://www.adobe.com/uk/ |
| Photoshop | Adobe | https://www.adobe.com/uk/ |
| MaxQuant | Nat Protoc. 2016 Dec;11(12):2301-2319. Version v1.6.10.43 | http://www.maxquant.org |

### Method details

#### Cell culture

THP-1 cells were cultured in 10% FBS (Gibco #10500-64) RPMI 1640 (Sigma) supplemented with 2% L-glutamine (G7513, Sigma-Aldrich), 0.05 mM β-mercaptoethanol (Sigma), and antibiotics (penicillin/streptomycin, Gibco). HEK293T cells were cultured in high glucose DMEM, supplemented with 10% FBS, and 2% L-glutamine. Flp-In T-REx 293 cell line (R78007, ThermoFisher) were cultured in high glucose DMEM, supplemented with 10% FBS, 2% L-glutamine, 100 µg/mL Zeocin (InvivoGen, ant-zn-1) and 15 µg/mL blasticidin (InvivoGen, ant-bl-1) at 37 °C in a humidified 5% CO2 atmosphere. The stably transfected Flp-In T-REx 293 cell lines (flag-NLRP3 and flag-NLRP3-APEX2) were cultured in the same medium and selected by 100 μg/mL hygromycin B (InvivoGen, ant-hg-1). 1 μg/mL tetracycline was added to induce the expression of flag-NLRP3 or flag-NLRP3-APEX2.

#### PBMC isolation and *in vitro* culture

To isolate the mononuclear cells, leukocyte cones (around 4 mL) were diluted in 25 mL RPMI medium before laid onto 15 mL LymphoPrep without disturbing the interface between LymphoPrep and the cone blood. Subsequently, it was centrifuged at 2000 rpm (400 g) at room temperature for 20 min. Lymphocytes, which form a thin layer of white film after the centrifuge, are transferred into a fresh 50 mL tube with a Graduated Pasteur Pipette and washed twice with RPMI medium. PBMCs were plated in 6-well plate at 1-2 × 10^7^ cells/mL in 10 mL and allowed to adhere at 37℃ for 4 hrs. Non-adherent cells were removed thorough washing with RPMI-1640.

#### hiPSC culture and differentiation to microglia

The KOLF2.1S hiPSC line used for this study was obtained with informed consent from all subjects, donors and/or their legal guardian(s) and with the approval of all the relevant institutions. All methods and experimental protocols were carried out in accordance with University of Oxford guidelines and regulations. hiPSC lines were generated previously as part of the HipSci project (REC Reference 09/H0304/77, covered under HMDMC 14/013 and REC reference 14/LO/0345). KOLF2.1S was generated from the parental KOLF2-C1 line (https://hpscreg.eu/cell-line/WTSIi018-B-1) itself a clonal isolate of KOLF2 (https://hpscreg.eu/cell-line/WTSIi018-B) by correction of a heterozygous loss of function mutation ARID2 using CRISPR based homology repair by the Cellular Operations Gene Editing team at the Wellcome Sanger Institute (Bruntraeger et al., 2019). hiPSC were cultured in OxE8 medium with daily media changes (Vaughan-Jackson et al., 2021) on Geltrex (Gibco A1413302) coated tissue culture dishes and passaged with 0.5mM EDTA. hiPSC were differentiated into microglial precursors through the formation of embryoid bodies (EB) (van Wilgenburg et al., 2013) by seeding 4 x 10^6^ hiPSC per well of an Aggrewell 800 plate (STEMCELL Technologies, 34815) in EB medium (OxE8 supplemented with 50 ng/mL BMP4 (PeproTech PHC9534), 50 ng/mL VEGF (PeproTech PHC9394), and 20 ng/mL SCF (Miltenyi Biotec 130-096-695)). After differentiating in EB media for 6 days (with daily 50% media changes), 150 EB were transferred to T175 coated with 0.1% Gelatin containing differentiation media (X-VIVO 15 (Lonza BE02-060F), 1x GlutaMAX (Lifetech 25050- 061), 50µM β-mercaptoethanol (LifeTech 31350-010), 100 ng/mL M-CSF (Gibco, PHC9501), 25 ng/mL IL-3 (Gibco, PHC0033). After 4-5 weeks of differentiation, microglial precursors shed into the media are harvested weekly by collection of supernatant and pelleting at 400 x g for 5 minutes. Microglial precursors are then resuspended in the final microglia maturation media ITMG (ADMEM/F12 (LifeTech, 12634-010), 1x GlutaMAX (Lifetech, 25050-061), 100 ng/mL IL-34 (PeproTech, 200-34-500), 50 ng/mL TGFβ-1 (PeproTech, 100-21C-250), 25 ng/mL M-CSF (Gibco, PHC0033), 10 ng/mL GM-CSF (Gibco, PHC2013), as previously described (Washer et al., 2022), and differentiated to hiPSC-microglia for 14 days, with 50% media changes every 3 to 4 days. All differentiation steps are carried out at 37°C, 5% CO2.

#### Inflammasome activation assay

THP-1 cells were plated in a concentration of 1x10^6^ cells/mL and incubated with 0.5 μM PMA for 4 h. After differentiation, PMA-containing media was replaced with fresh media. Experiments were carried out on the following day. Differentiated THP-1 cells were primed with LPS (1 µg/mL) for 4 h, followed by the treatment of PBS (mock) or 5 μM nigericin for 45 min. After stimulation, supernatants and cell lysates were collected separately, or combined together for immunoblotting analysis.

#### ASC speck oligomer crosslinking

PMA-differentiated THP-1 cells were primed with LPS (1 µg/mL) for 4 h, and then stimulated with nigericin (10μM) for 45 min. Cells were lysed on ice in Buffer A (20 mM HEPES-KOH, pH 7.5, 10 mM KCl, 1.5 mM MgCl_2_, 1 mM EDTA, 1 mM EGTA, 320 mM sucrose, and protease inhibitor cocktail), and homogenized using a 21-gauge needle. The lysates were centrifuged at 1,800 rpm (300 g) for 8 min at 4 ℃. Supernatants were transferred to a new tube and diluted with 1 volume of CHAPS buffer (20 mM HEPES-KOH, pH 7.5, 5 mM MgCl2, 0.5 mM EGTA, 0.1% CHAPS, and protease inhibitor cocktail). The diluted lysates were then centrifuged at 5,000 rpm (2,400 g) provide g for 8 min to obtain the crude inflammasome-containing pellets. The pellets were washed with ice cold PBS twice, resuspended in CHAPS buffer, and then cross-linked at 37 ℃ for 20 min by adding 2mM DSS (disuccinimidyl suberate, dissolved in DMSO). The reaction was quenched by adding SDS sample buffer and followed by Western blotting analysis.

#### Enzyme-linked immunosorbent assay (ELISA)

Human IL-1β (DY201, R&D Systems) and IL-6 (DY206, R&D Systems) production in the supernatants were detected by ELISA kit according to manufacturer’s protocol. Plates were coated with diluted capture antibody in sterile PBS at room temperature overnight. On the following day, plates were washed three times with 200 μL/well wash buffer (0.05% Tween 20 in PBS, pH 7.2-7.4) before being blocked with 200 μL/well reagent diluent (1% BSA in PBS, pH 7.2-7.4, 0.2 μm filtered) at room temperature for a minimum of 1 h. After blocking, plates were washed three times and incubated with samples or standards (in reagent diluent) for 2 h at room temperature. After three washes of plates, 100 μL/well detection antibody (diluted in reagent diluent) was added to the wells and incubated for 2 h at room temperature. Plates were then washed three times and incubated with 100 μL/well Streptavidin-HRP for 20 min at room temperature, followed by another three washes and incubation with 100 μL/well substrate solution (1:1 mixture of Colour Reagent A (H_2_O_2_) and Colour Reagent B (Tetramethylbenzidine)) at room temperature. 20 min after the reaction, 50 μL/well stop solution (2 N H2SO4, DY994, R&D Systems) was added and mixed thoroughly to quench the reaction. The optical density of each well was determined at the wavelength of 450nm, and the relative concentration was calculated based on the standard curve, generated through a 3 parameter logistic curve fit using the data from standard samples.

#### TCA/DOC protein precipitation

Trichloroacetic acid (TCA)- deoxycholate (DOC) precipitation was commonly used to concentrate the proteins in supernatants and this method was adapted from André et al. Briefly, 8.5 μL of 2% Na-deoxycholate was added to 1 mL sample with a final concentration of 125 μg/mL. This mixture was incubated at room temperature for 15 min and 333 μL of 24% TCA (final concentration 6%) was added to the samples. Proteins were precipitated and centrifuged at 12,000 g for 30 min at 4°C. Supernatant was carefully removed and the pellets were washed once with acetone to remove excessive TCA before another centrifuge at 12,000 g for 5 min. After removing the supernatants, the resulting pellets were solubilised in loading buffer and subjected to immunoblotting analysis.

#### Cytotoxicity and cell viability assay

Relevant cells were treated as indicated, and the cytotoxicity was detected using CyQUANT™ LDH Cytotoxicity Assay (Invitrogen) according to manufacturer’s instruction. Briefly, 50 μL of each sample supernatant was transferred to 96-well flat-bottom plate in triplicate wells, and then, 50 μL of premixed reaction buffer was added to each well. The plate was incubated for 30 min at room temperature before addition of 50 μL/well stop solution. The optical density of each well was measured at the wavelength of 490nm and 680nm (680 nm absorbance was used as background), and the cytotoxicity (% LDH release) was calculated according to manufacturer’s protocol.

Cell viability was measured using CellTiter 96 AQueous One Solution Cell Proliferation Assay (MTS, Promega). Briefly, 20µl of CellTiter 96® AQueous One Solution Reagent was added to each well containing samples in 100 µl of culture medium. The plate was then incubated for a minimum of 1h at 37°C. The absorbance of each well was measured at the wavelength of 490nm, and the cell viability was calculated according to manufacturer’s protocol.

#### Caspase-1 activity bioluminescent assay

Relevant cells were treated as indicated, and the caspase-1 activity in the culture medium (supernatant) was measured using Caspase-Glo® 1 Inflammasome Assay (Promega) according to manufacturer’s instruction. Briefly, Caspase-Glo 1 Reagent was prepared through resuspension of Z-WEHD substrate using Caspase-Glo 1 buffer, and the addition of MG132 inhibitor to a final concentration of 60 μM. Afterwards, 50 μL of cell culture medium was transferred to the 96-well plate, followed by adding 5 μL Caspase-Glo 1 Reagent. The contents of the wells were thoroughly mixed and incubated at room temperature for at least 1 hour for the signal to be stable. Subsequently, the luminescence value was measured using multi-mode microplate reader FLUOstar Omega (BMG LABTECH) according to manufacturer’s instruction.

#### Immunoprecipitation

Cells were washed with ice-cold PBS for two times, and then lysed in Pierce® IP lysis buffer (25 mM Tris-HCl pH 7.4, 150 mM NaCl, 1% NP-40, 1 mM EDTA, 5% glycerol, PhosSTOP phosphatase inhibitor and cOmplete™ Protease Inhibitor Cocktail). Lysates were incubated on ice for 10 min with periodic mixing, and then clarified at 13,000 g for 10 min at 4°C to remove the cell debris. Pre-cleared lysates were incubated with anti-FLAG® M2 Magnetic Beads at room temperature for 30 min. After three times of washes with TBS, the proteins bound by the anti-flag antibody were pulled down and 3X FLAG® peptide was added for elution. Both the whole cell lysates and the pull-down samples are subjected to immunoblotting analysis.

#### Transient transfection

For DNA plasmids: Cells were grown in 6-well plates and Lipofectamine 3000 Transfection Reagent (L3000001, ThermoFisher UK) was used to transfect the cells following the manufacturer’s instructions. The concentration of plasmids used was indicated in each experiment. For siRNA transfection experiments: Cells were grown in 6-well plates and RNAimax transfection reagent (Invitrogen, 13778- 150) was used following the manufacturer’s protocol. A final concentration of 10 nM of the following Silencer™ select siRNAs were used: control siRNA (4390844), si-NLRP3 (s41554) and si-UCH-L1 (s14616).

#### Construction of plasmids

The full list and descriptions of all primers and plasmids used in this study can be found in key resources table. For the generation of NLRP3-APEX2, NLRP3-Flag, NEBuilder HiFi DNA Assembly (NEB: E5520S) was used following manufacture protocol. Primers were designed using the NEBuilder Assembly Tool. The successful clones were confirmed using both PCR and DNA sequencing. Finally, the expression plasmids for NLRP3 domains were generated using the Q5® Site-Directed Mutagenesis Kit (NEB: E0554S) following manufacture protocols. We used as a template the GFP- NLRP3 plasmid (Addgene), we initially replace GFP with Flag. Finally, different domains were deleted to generate the successful plasmids. The successful clones were confirmed using both PCR and DNA sequencing.

#### Generation of stable cell line

The Flp-In T-Rex 293 stable cell lines (flag-NLRP3 and flag-NLRP3-APEX2) were generated using Lipofectamine 3000 following manufacture protocol in a 6-well plate. The cells were transfected with both the expression plasmid and the plasmid pOG44 in a ratio of 1:9. Cells were split the following day and transferred in a 100mm dish. After 8 hours, 300 μg/mL hygromycin B was added into the medium for selection.

#### sgRNA design, lentivirus production and transduction

Non-targeting control (NTC) and UCH-L1-targeting sgRNA sequences were designed using CRISPick tool (https://portals.broadinstitute.org/gppx/crispick/public). To clone the target sequences into the lentiCRISPRv2 backbone (one vector system), two oligos containing the target sequences were synthesized (Sigma) and inserted into the digested lentiviral construct. sgRNA sequences are as followed: UCH-L1-sgRNA#1: 5’-ACTTCATGAAGCAGACCATT-3’; UCH-L1-sgRNA#2: 5’-AATCGGACTTATTCACGCAG-3’; UCH-L1-sgRNA#3: 5’- AAGACAAACTGGGATTTGGT-3’.

HEK293T cell lines were seeded at 40% confluency and transfected with pMD2G, psPAX2 and the above lentiCRISPRv2 plasmids using lipofectamine 2000 following manufacturer’s instructions. After incubation for two days, supernatants were collected and filtered by 0.45µm filter, followed by ultracentrifuge to obtain concentrated viral particles.

#### Biotin-phenol proximity labelling and biotinylated proteins capture

This method was adapted from that previously reported by Stephanie et al. 5×10^6^ Flp-In T-Rex HEK293 stable cells were induced with 1 μg/mL tetracycline overnight. The next day cells were incubated with medium containing 500 μM biotin-phenol (BP) for 30 min at 37 °C under 5% CO2. Afterwards, a final concentration of 1 mM H_2_O_2_ was added and incubated for 40 seconds. The reaction was then quenched by the addition of quenching buffer (10mM sodium ascorbate, 10mM sodium azide, 5mM Trolox in DPBS). Subsequently, cells were washed with quenching buffer for three times, PBS for two times and then lysed with RIPA lysis buffer containing 1 mM PMSF, 5 mM Trolox, 10 mM sodium azide, 10 mM sodium ascorbate, PhosSTOP phosphatase inhibitor and cOmplete™ Protease Inhibitor Cocktail. Samples were then briefly sonicated and centrifuged at 10,000 g for 10 min at 4°C to remove cell debris. For streptavidin pull-down experiments, Dynabeads MyOne Streptavidin T1 was prepared and washed with RIPA lysis buffer twice. The cleared lysate was applied to the magnetic beads and at RT for 1 hour. Afterwards, the beads were washed for two times with RIPA lysis buffer, once with 1M KCL, 0.1 M Na_2_CO_3_ and 2 M Urea (in 10 mM Tris-HCL, pH 8) and finally twice with RIPA buffer again. For western blot analysis, biotinylated proteins were eluted using 60 μL of 3×loading buffer supplemented with 2 mM biotin and 20 mM DTT by boiling (98°C) for 10 min. For Mass spectrometry analysis, an on-beads digestion protocol was used as described below. Samples were denatured in 8 M Urea (in 100 mM triethylammonium bicarbonate buffer (TEAB)) and incubated at room temperature for 30 min. Afterwards, the samples were reduced with 10mM tris(2-carboxyethyl)phosphine (TCEP) for at room temperature 30 min. Samples were alkylated by the addition of iodoacetamide (final concentration 50 mM) and incubated at room temperature for 30 min. The reaction was quenched by adding DTT (final concentration 20 mM), and the urea concentration were diluted down to 1.5 M by adding TEAB. Biotinylated proteins were digested through the incubation with trypsin (1 μg trypsin per 20 μL streptavidin beads) overnight at 37 ℃. Following the trypsinization, samples were desalted using the C18 solid phase cartridge (Sep-Pak, Waters) according to the manufacturer’s protocol. Purified peptide eluates were dried down in the speed-vac, re-suspended in buffer A (98 % MilliQ-H20, 2 % CH_3_CN and 0.1 % trifluoroacetic acid - TFA) and stored at -20 °C until further analysis.

#### Affinity-purification mass spectrometry (AP-MS)

1×10^7^ HEK293 FLAG-NLRP3 Flp-In T-Rex stable cells were induced with 1 μg/mL tetracycline overnight. The next day cells were incubated with medium with Nigericin (10μM) or DMSO for 90 min at 37 °C under 5% CO_2_. Cells were washed with ice-cold PBS for five times, and then lysed in Pierce IP lysis buffer (25 mM Tris-HCl pH 7.4, 150 mM NaCl, 1% NP-40, 1 mM EDTA, 5% glycerol, PhosSTOP phosphatase inhibitor and cOmplete™ Protease Inhibitor Cocktail). Lysates were incubated on ice for 10 min with periodic mixing, and then clarified at 13,000 g for 10 min at 4°C to remove the cell debris. Pre-cleared lysates were incubated with anti-FLAG M2 Magnetic Beads at 4°C overnight. The next day beads were washed 3 times with TBS. Samples were eluted were eluted using 100 μL of 3×loading buffer supplemented 20 mM DTT by boiling (98°C) for 10 min.

#### Mass spectrometry analysis

An Orbitrap Fusion Lumos Tribid mass spectrometer (Thermo Fisher Scientific) coupled to an Ultimate 3000 UHPLC (Thermo Fischer Scientific) platform was used to analyse the purified tryptic peptides by liquid chromatography tandem mass spectrometry (LC-MS/MS) as described previously (Davis et al., 2017). In brief, ten per cent of tryptic peptides (∼ 200 ng) were loaded onto a trap column (PepMapC18; 300µm x 5mm, 5µm particle size, Thermo Fischer) and separated on a 50 cm EasySpray column (ES803, Thermo Fischer) using a 60 minute linear gradient ranging from 2 % to 35% buffer B (A: 5% DMSO, 0.1% formic acid; B: 5% DMSO, 0.1% formic acid in acetonitrile) at a flow rate of ∼250 nL/min. Eluted peptides were then subjected to electrospray ionisation (ESI) and directed into an Orbitrap Fusion Lumos Tribrid mass spectrometer (instrument control software v3.3). Data were acquired in data-dependent mode (DDA), with the advance peak detection (APD) feature enabled. Survey scans were acquired in the Orbitrap at 120 k resolution over a m/z range 400 -1500, AGC target of 4e5 and S-lens RF of 30. Fragment ion spectra (MS/MS) were obtained in the Ion trap (rapid scan mode) with a Quad isolation window of 1.6, 40% AGC target and a maximum injection time of 35 ms, with HCD activation and 28% collision energy. Raw MS files were analysed using MaxQuant software (v1.6.10.43) using label free quantitation (LFQ) as described previously (Olie et al., 2023). In brief, raw MS files were searched against the UniProtKB human sequence database (UP000005640, 79,038 human sequence entries). The following parameters were set to perform label free quantification: Modifications were set on Carbamidomethyl (C) as fixed and Oxidation (M) and Deamidation (NQ) as variable and a maximum of two missed cleavages were allowed. Match between runs function was used.

#### RNA extraction and real time reverse transcription (RT)-PCR

Total RNA was isolated with the TRIzol reagent (SigmaAldrich) according to the manufacturer’s instructions. The nucleic acid concentration was measured with the NanoDrop2000 (NanoDrop Technologies, Inc.). Complementary DNA (cDNA) was synthesized from 1.5 µg of total RNA with the High-Capacity cDNA Reverse Transcription Kit (Applied Biosystems, USA). Quantitative real-time polymerase chain reaction (RT-qPCR) was performed using the LightCycler 480 SYBR Green I Master kit and LightCycler 480 PCR System (Roche, Switzerland) according to the manufacturer’s instructions. The Delta-Ct method was used to calculate the relative RNA expression levels.

#### Pathway enrichment analysis

Filtered proximity proteomes (SAINT probability > 0.8, FC-A ≥ 1) either in DMSO control or nigericin activated samples were imported into the STRING database search portal (version 11.5, Szklarczyk et al., 2021; von Mering et al., 2003), and the top 20 retrieved terms were plotted with both strength (Log10 [observed / expected]) and false discovery rate (FDR) (interaction score ≥ 0.7). Terms were clustered through the similarity of enrichment strength patterns across the four groups and the results were further visualized using clusterProfiler 4.0 (Wu et al., 2021; Yu et al., 2012).

#### PPIs network construction

Differentially enriched proteins under unstimulated or stimulated condition that passed the filtering criteria (log2[FC]≥0.5 or ≤-0.5, FDR≤ 0.05) were separately uploaded to the STRING portal, and physical subnetwork type was chosen to extract the physical (direct) protein-protein associations from the databases. Active interaction sources, including text mining, experiments, and databases, and an interaction score > 0.7 (high confidence interaction) were applied to construct the PPI networks. Markov clustering algorithm (MCL) was used to group the interaction subnetworks with the inflation value of 3.0, resulting in 13 clusters for the stimulated condition and 18 clusters for the unstimulated condition.

#### SAINT probabilistic scoring for NLRP3 APEX2 PL-MS and AP-MS data

The computational tool, SAINTexpress (Teo et al., 2014)- an updated version of the “Significance Analysis of INTeractome (SAINT),” was utilised to allocate confidence scores to the proximity labelling and AP-MS data. This analysis was conducted via the online Contaminant Repository for Affinity Purification platform at www.crapome.org (Mellacheruvu et al., 2013).

### Statistics and reproducibility

Data are presented as mean ± standard error of the mean (SEM). Two-tailed, unpaired student’s t test was used for statistical analysis in GraphPad Prism 9. Unless stated, otherwise all quantitative experiments were performed in triplicates and average with SEM reported.

## Data availability

The mass spectrometry proteomics data have been deposited to the ProteomeXchange Consortium via the PRIDE (Deutsch et al., 2023; Perez-Riverol et al., 2022) partner repository with the dataset identifier PXD045862.

## Acknowledgements

Z.L., A.D., I.V., G.L., X.Z., R.F., T.D. and B.M.K. were supported by the Chinese Academy of Medical Sciences (CAMS) Innovation Fund for Medical Science (CIFMS), China (grant number: 2018-I2M-2- 002) awarded to B.M.K., R.F. and T.D. B.M.K. was supported by Pfizer. E.D. was supported by Alzheimer’s Research UK Grants ARUK2015DDI-OX. E.J was funded by Wellcome Trust (grant # 224040/Z/21/Z). Imaging was performed using the Oxford-Zeiss Centre of Excellent in Biomedical Imaging and the Kennedy Trust for Rheumatology Research (grant # 202117 and 202103, respectively).

E.W.T. was supported by a CRUK Programme Award (DRCNPG-Nov21\100001) with support from the Engineering & Physical Sciences Research Council. We would also like to thank members of the Discovery Proteomics Facility, the Kessler, Oxford Drug Discovery Institute groups for constructive discussions and Raphael Heilig for his expert help with mass spectrometry.

## Author contributions

B.M.K., E.D. and Z.L. designed the research and wrote the manuscript. Z.L. and A.D. performed the NLRP3-APEX2 proximity labelling experiments and analysed the mass spectrometry data. Z.L. and A.D. performed the DUB siRNA library screen experiment. I.V. operated mass spectrometer for LC-MS/MS acquisition of the samples and contributed to data processing. E.J. performed confocal imaging and analysis. S.J.W. and A.D. contributed to the culture of hiPSC and differentiated them into microglial cells. E.W.T. and F.C. contributed to UCH-L1 inhibitor provision and study design. Z.L., A.D., I.V., E.J., F.H.L., S.J.W., G.L., G.Y., H.L., F.C., X.Z., R.F., T.D., E.W.T., E.D. and B.M.K discussed the results, commented, and provided critical reviews on the manuscript.

---

**Figure S1.**
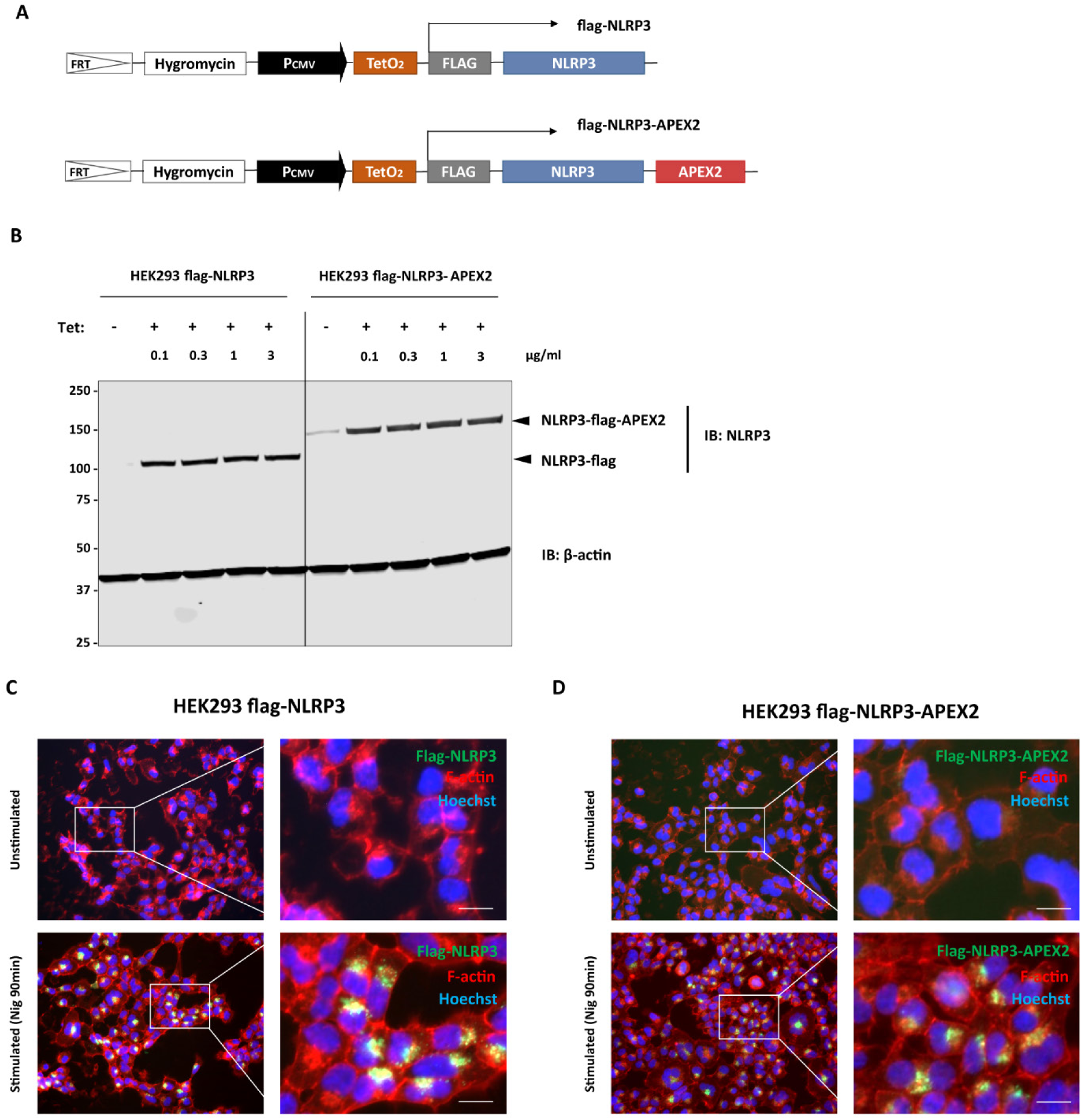
Construction of flag-NLRP3 and flag-NLRP3-APEX2 HEK293 stable cell line. (A) Schematic of Flag-NLRP3 and Flag-NLRP3-APEX2 constructs. (B) Both the flag-NLRP3 and flag- NLRP3-APEX2 Flp-In T-Rex HEK293 stable cell lines were treated with tetracycline at indicated concentrations (0.1-3 μg/mL). 24 hours after induction, cell lysates were collected for immunoblotting analysis. (C-D) Immunofluorescence of flag-NLRP3 (C) and flag-NLRP3-APEX2 (D) in Flp-In T-Rex HEK293 stable cell lines. Cells were either treated with DMSO or Nigericin (10μM) for 90 min. Scale bars, 10 μm.

**Figure S2.**
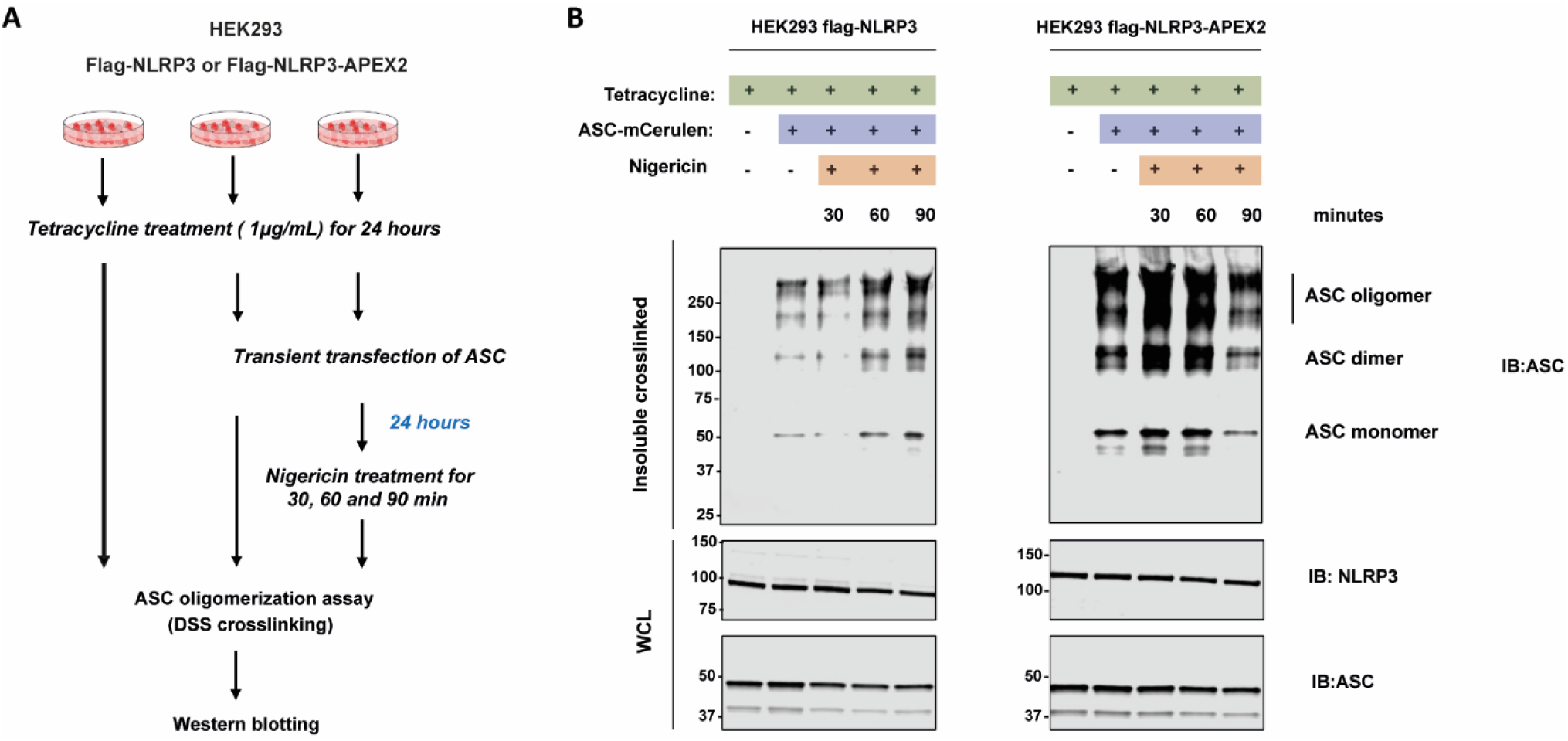
Reconstituted NLRP3 induces ASC oligomerization in HEK293 cell lines. (A) Schematic of the ASC oligomerization assay in Flp-In T-Rex HEK293 stable cell lines. (B) Immunoblot of DSS-crosslinked ASC oligomers in the lysates from flag-NLRP3 and flag-NLRP3- APEX2 HEK293 stable cell lines, treated with tetracycline overnight, transfected with ASC mCerulean, and followed by the treatment with nigericin (10μM) for the indicated time.

**Figure S3.**
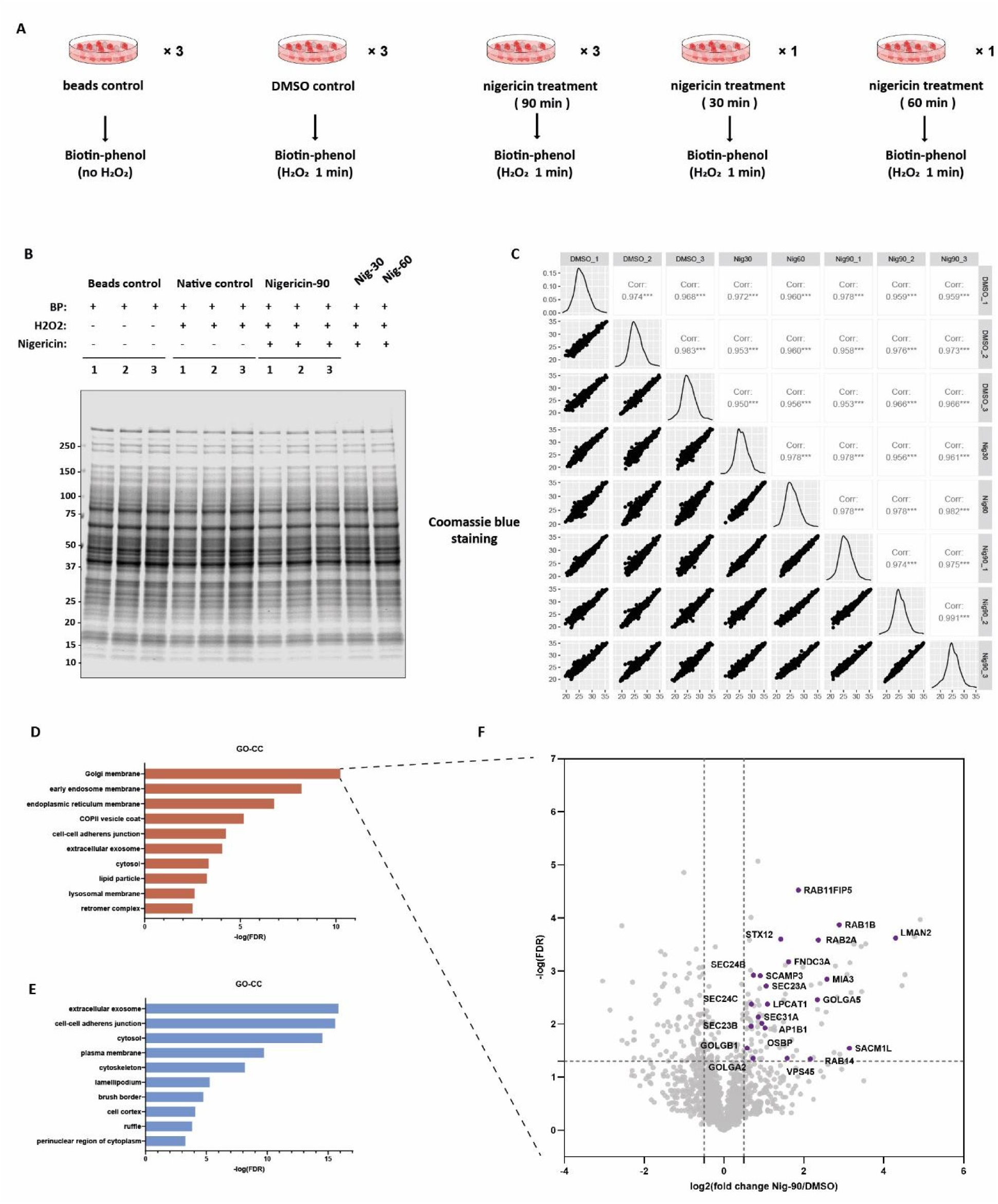
Tracing NLRP3 translocation using NLRP3-APEX2 proximity labelling. (A) Scheme of the experimental design and procedures of NLRP3-APEX2 proximity labelling. (B) Coomassie blue staining of whole-cell lysates from HEK293 Flp-In T-Rex NLRP3-APEX2 cells, treated as indicated with tetracycline, biotin-phenol, H_2_O_2_, and nigericin. (C) Pearson correlation matrix heatmap of the biotinylated proteome profiling. This analysis confirms high reproducibility in protein relative abundance measurements. (D-F) Bar graph showing the top 10 gene ontology-cellular component (GO-CC) terms of the proteins enriched in nigericin-stimulated state (nigericin treatment for 90 min) (D) and unstimulated state (DMSO treatment for 90 min) (E) respectively. (F) The 24 protein that belong to the Golgi membranes term are shown and coloured as purple in the volcano plotting.

**Figure S4.**
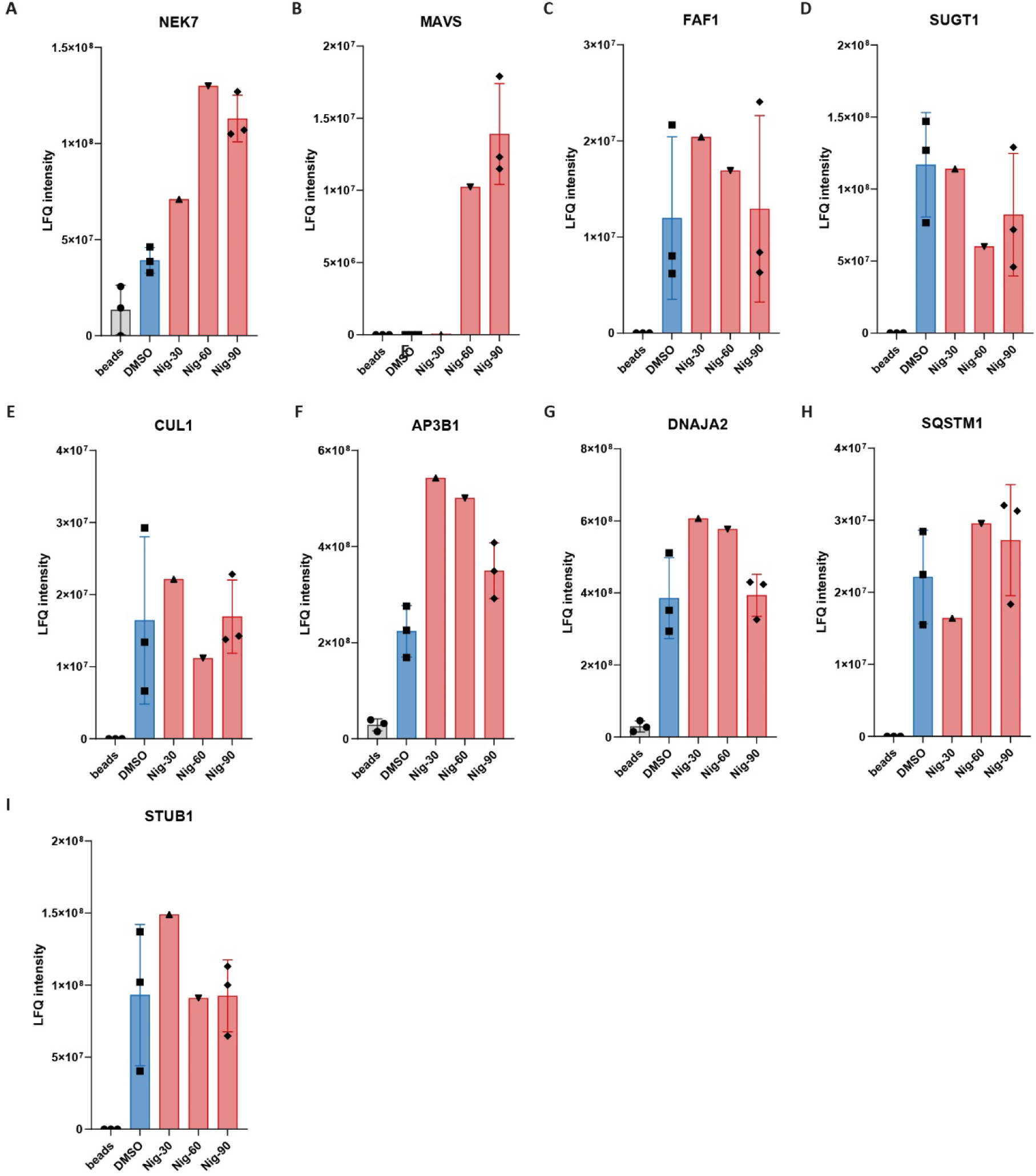
APEX2 proximity labelling identifies known NLRP3-interacting proteins during NLRP3 activation downstream of potassium efflux. (A-I) The raw LFQ Intensity values of the APEX2 NLRP3 proximity labelling from all the known NLRP3-interacting proteins identified in our experiment. An average value of 3 independent experiments is presented for DMSO and Nig-90 whereas for Nig-30 and Nig-60 just a single experiment. Error bars show the standard deviation of the mean.

**Figure S5.**
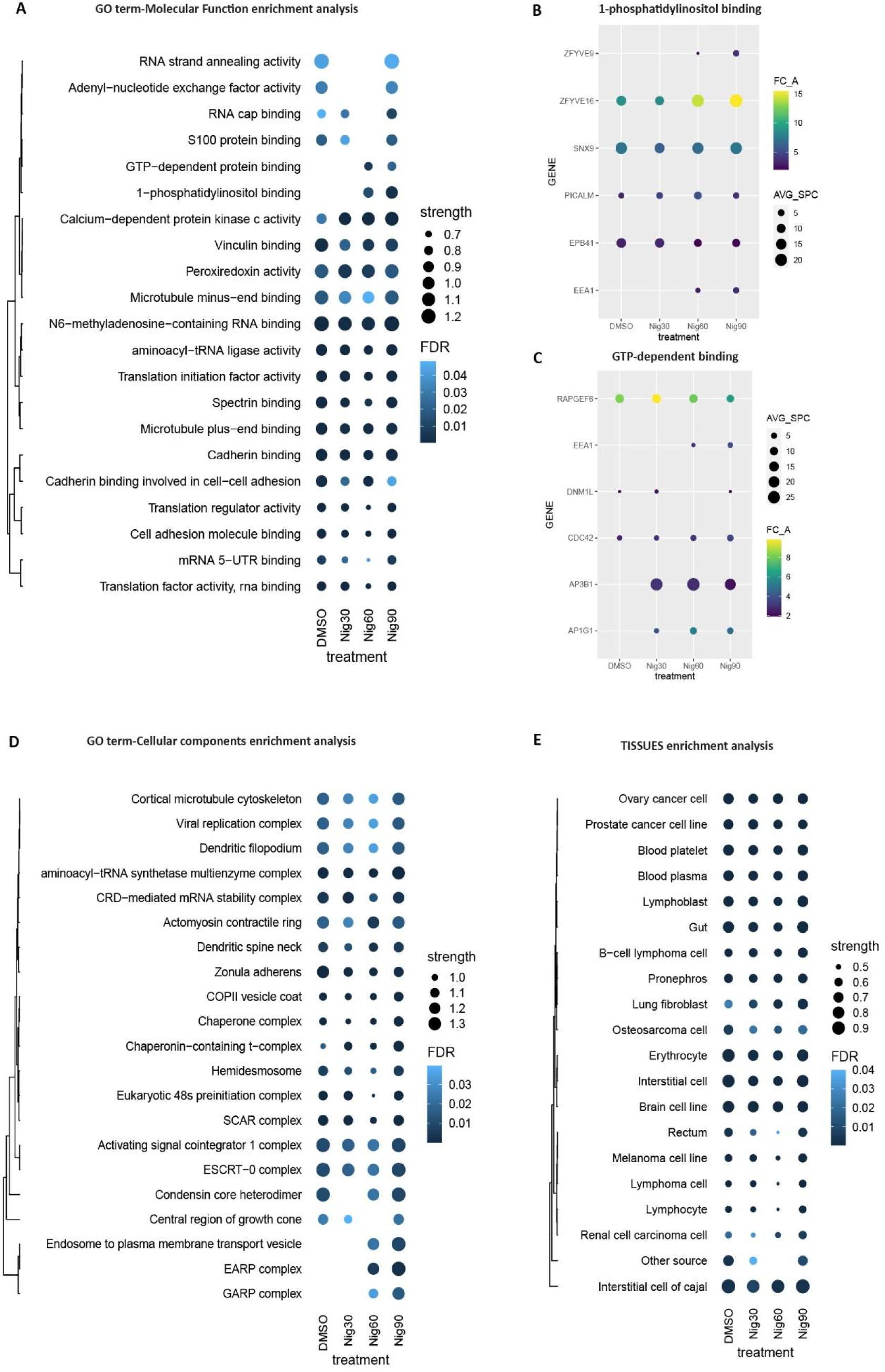
Functional analysis of the NLRP3 proximal proteins during inflammasome activation. (A, D, E) Dot plots displaying the GOBP (A), GOCC (D) and TISSUE (E) enrichment analysis of high confidence NLRP3 proximal proteins upon nigericin stimulation for different timepoints as indicated. (B, C) Dot plots showing the average spectral counts (AVG_SPC) and fold change (FC_A) of NLRP3 proximal proteins enriched in the GOMF term 1-phosphatidylinostol binding (B) and GTP-dependent binding (C) respectively.

**Figure S6.**
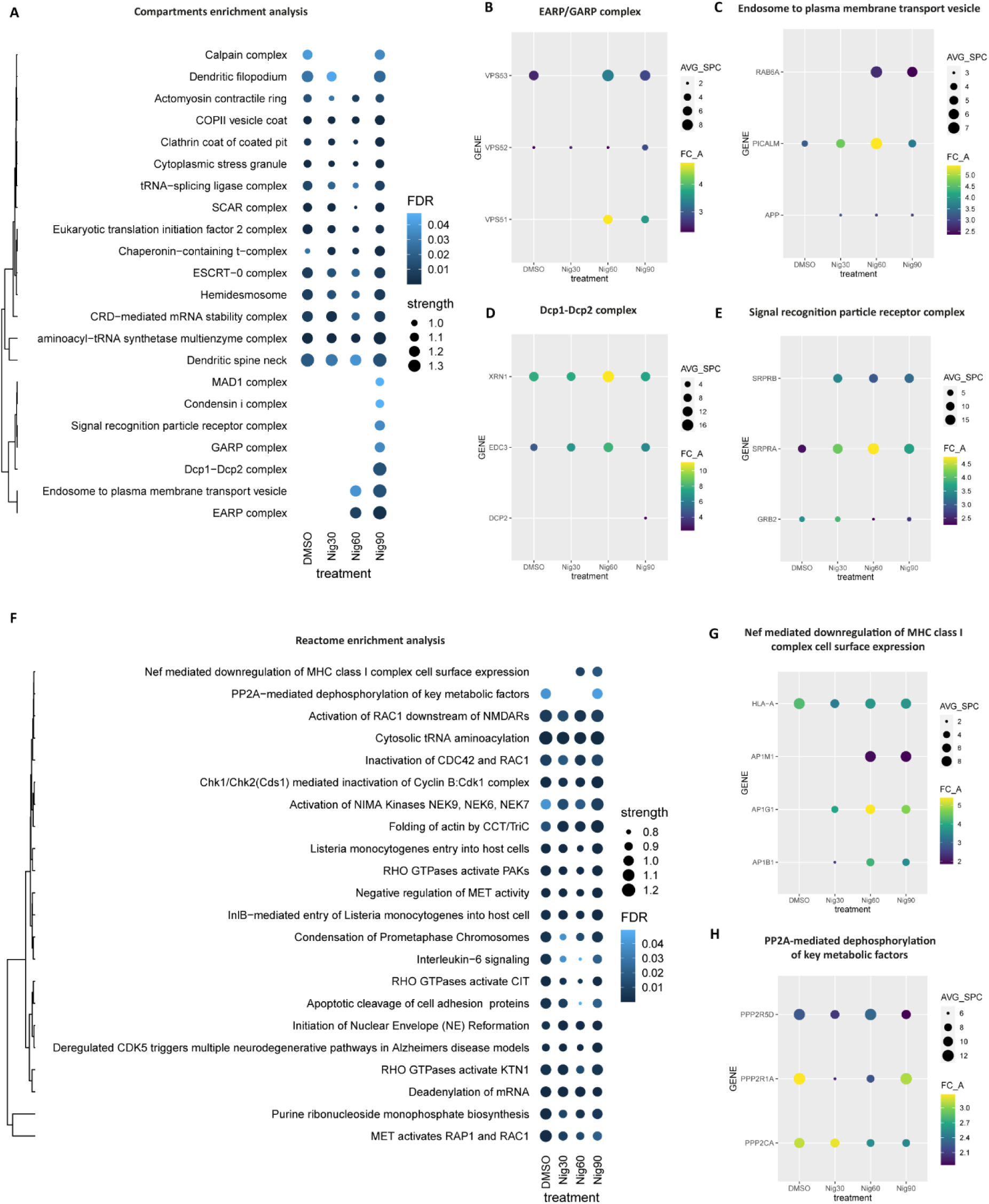
Functional analysis of the NLRP3 proximal proteins during inflammasome activation. (A, D) Dot plots displaying the Compartments (A) and Reactome (F) enrichment analysis of high confidence NLRP3 proximal proteins upon nigericin stimulation for different timepoints. (B-E, G-H) Dot plots showing the average spectral counts (AVG_SPC) and fold change (FC_A) of NLRP3 proximal proteins enriched in the term EARP/GARP complex (B), Endosome to plasma membrane transport vesicle (C), Dcp1-Dcp2 complex (D) and Signal recognition particle receptor complex (E), Nef mediated downregulation of MHC class I complex cell surface expression (G), PP2A-mediated dephosphorylation of key metabolic factors (H) respectively.

**Figure S7.**
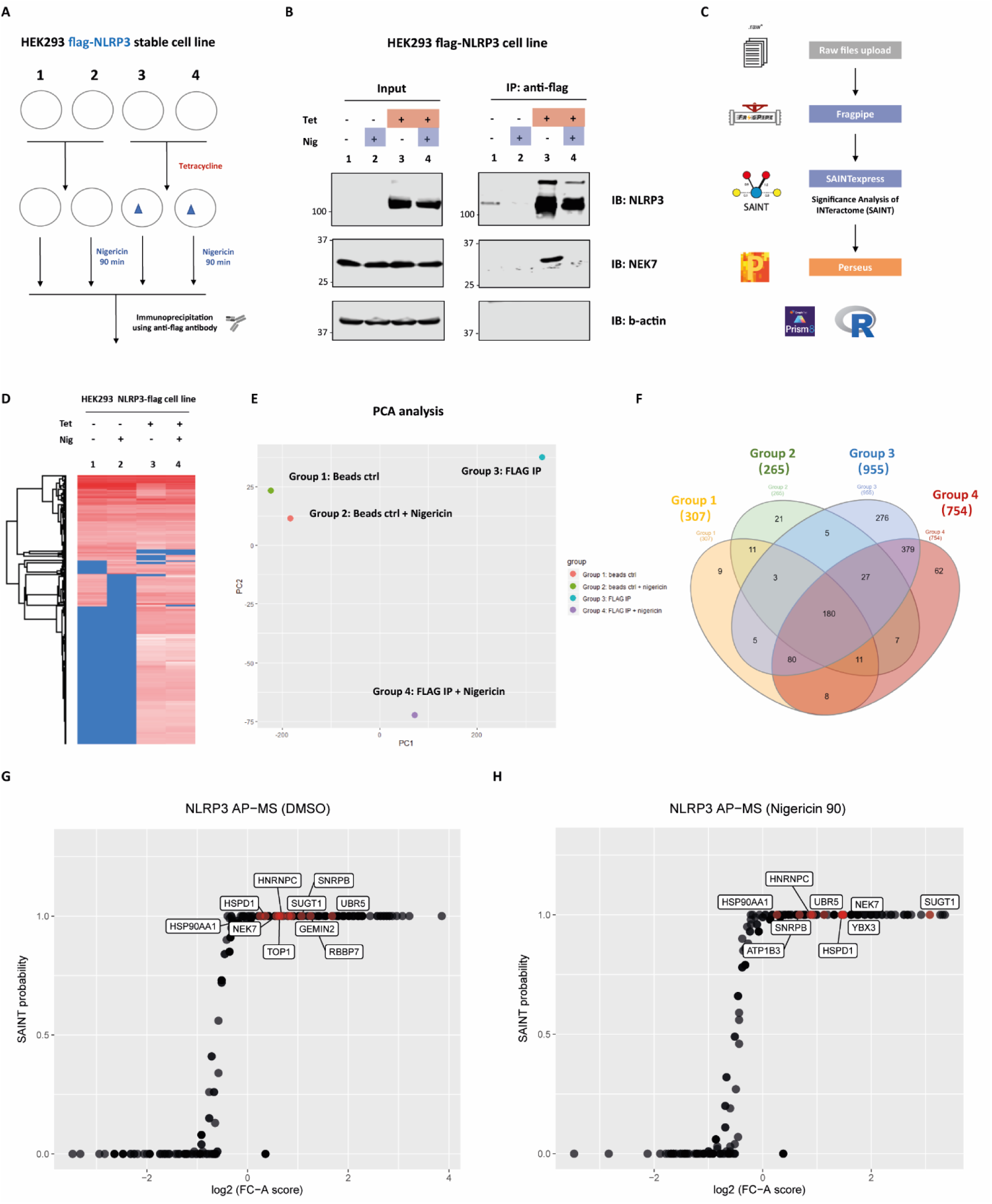
NLRP3 interactome determined by AP-MS. (A) Schematic of the NLRP3 AP-MS experiment design. (B) Stable HEK293 cells expressing Flag- NLRP3 were incubated with tetracycline overnight. Subsequently, they were treated with nigericin for 90 min as specified. NLRP3-flag was then isolated using an anti-flag antibody and subjected to immunoblotting with the indicated antibodies. (C) Diagram illustrating the data analysis workflow for AP-MS data. (D) Heatmap showing the hierarchical clusters of prey proteins in the NLRP3 AP-MS across the four groups. (E) Principal component analysis of the four samples. Each dot represents an individual sample. (F) Venn diagram illustrating the number of proteins identified from different groups as indicated in (A). (G-H) SAINT probability scoring of the proteomic data for NLRP3 AP-MS compared to the negative beads control. Proteins passing the cut-off (SAINT probability ≥ 0.8, FC-A ≥ 1) are considered as high-confidence interacting proteins for NLRP3.

**Figure S8.**
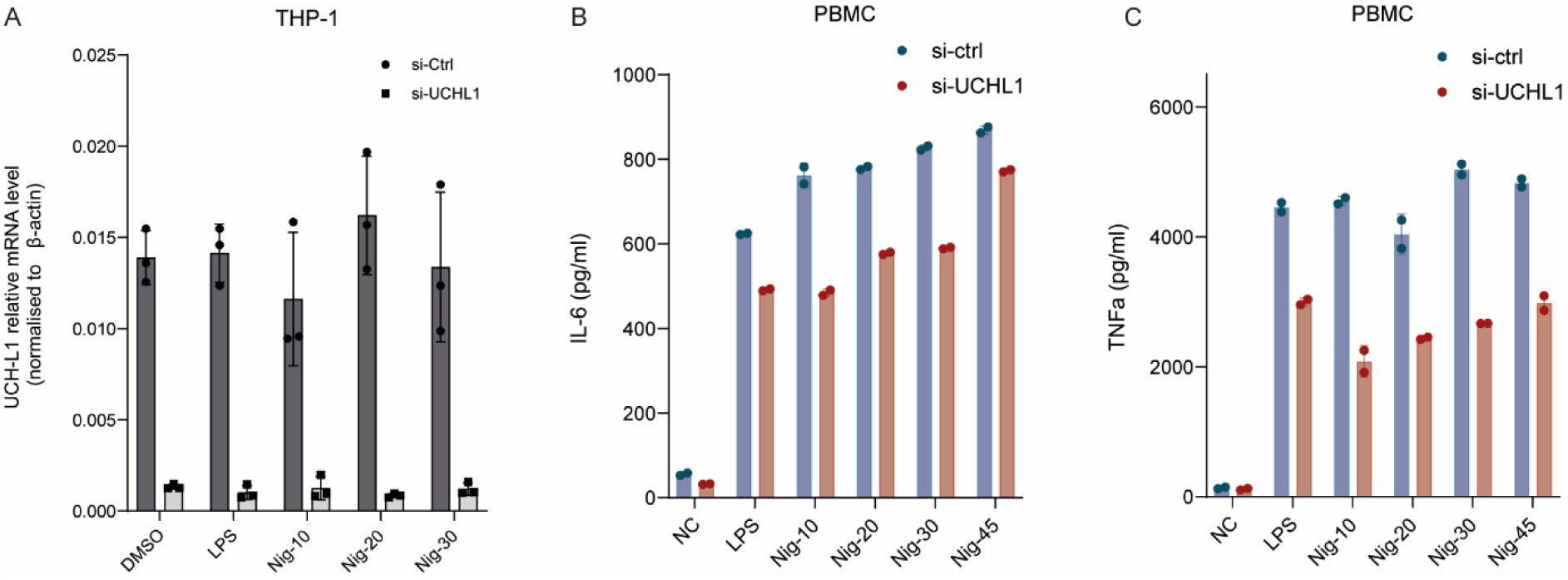
UCH-L1 deficiency decreases IL-6 and TNFα expression in human PBMC. (A) RT-qPCR analysis of UCH-L1 expression in THP-1 cells after transfection with either non-targeting siRNA or UCH-L1-targeted siRNA. Following differentiation with PMA, cells were primed using LPS (1 μg/mL) for 4 hours, and then treated with nigericin (10 μM) for different timepoints as indicated. (B-C) PBMCs were transfected with non-targeting control, or UCH-L1-specific siRNA, stimulated with LPS for 4 hours and nigericin for different timepoints as indicated. Production of IL-6 (B) and TNFα (C) in the supernatants from PBMCs were measured by ELISA.

**Figure S9.**
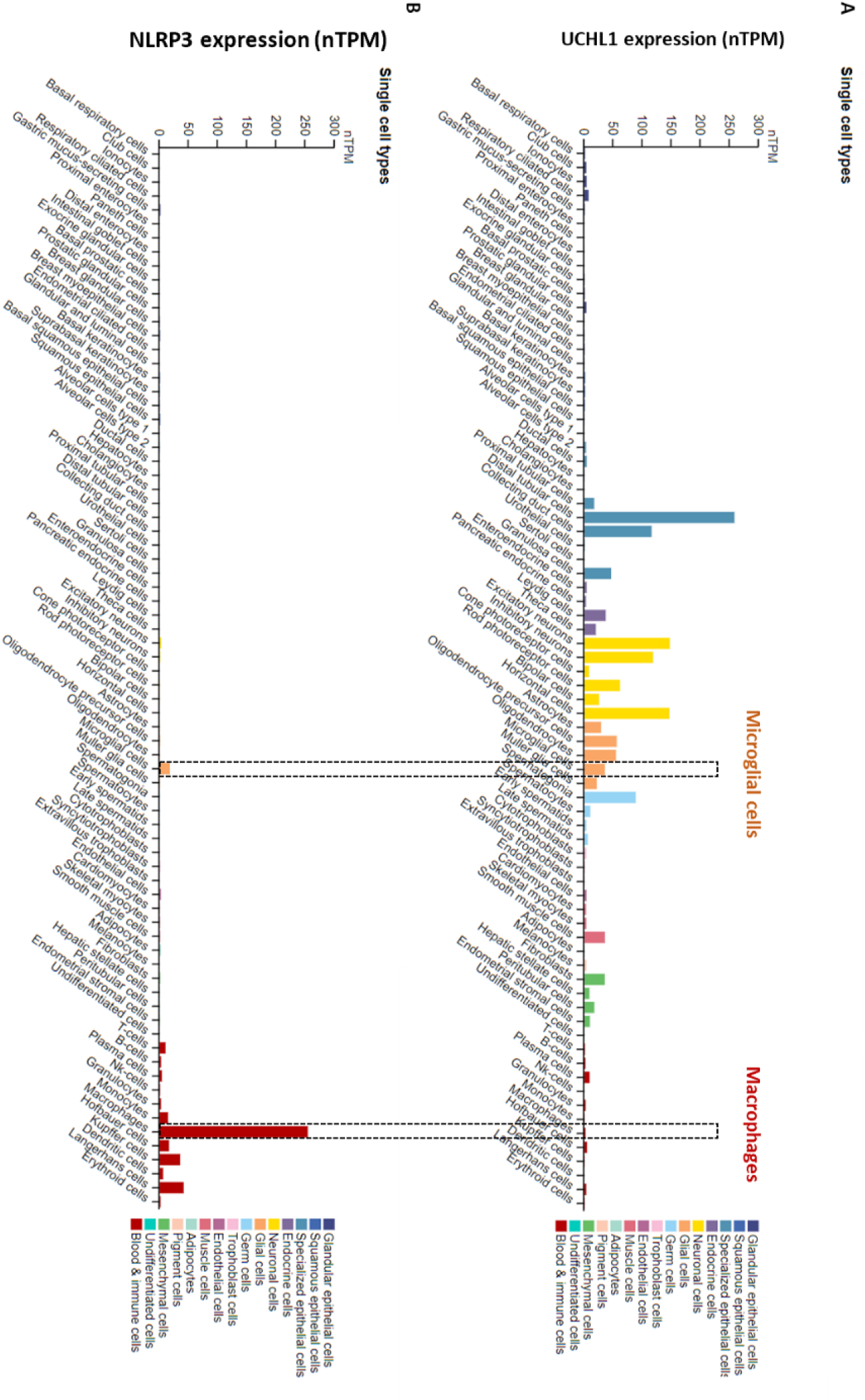
RNA expression level of NLRP3 and UCH-L1 in different cell types. (A,B) A summary of normalized single cell RNA expression (nTPM) in different cell lines for UCH- L1 (A) and NLRP3 (B). Color-coding represents different cell type groups consisting of cell types with functional features in common (Data were retrieved from https://www.proteinatlas.org/)

**Figure S10.**
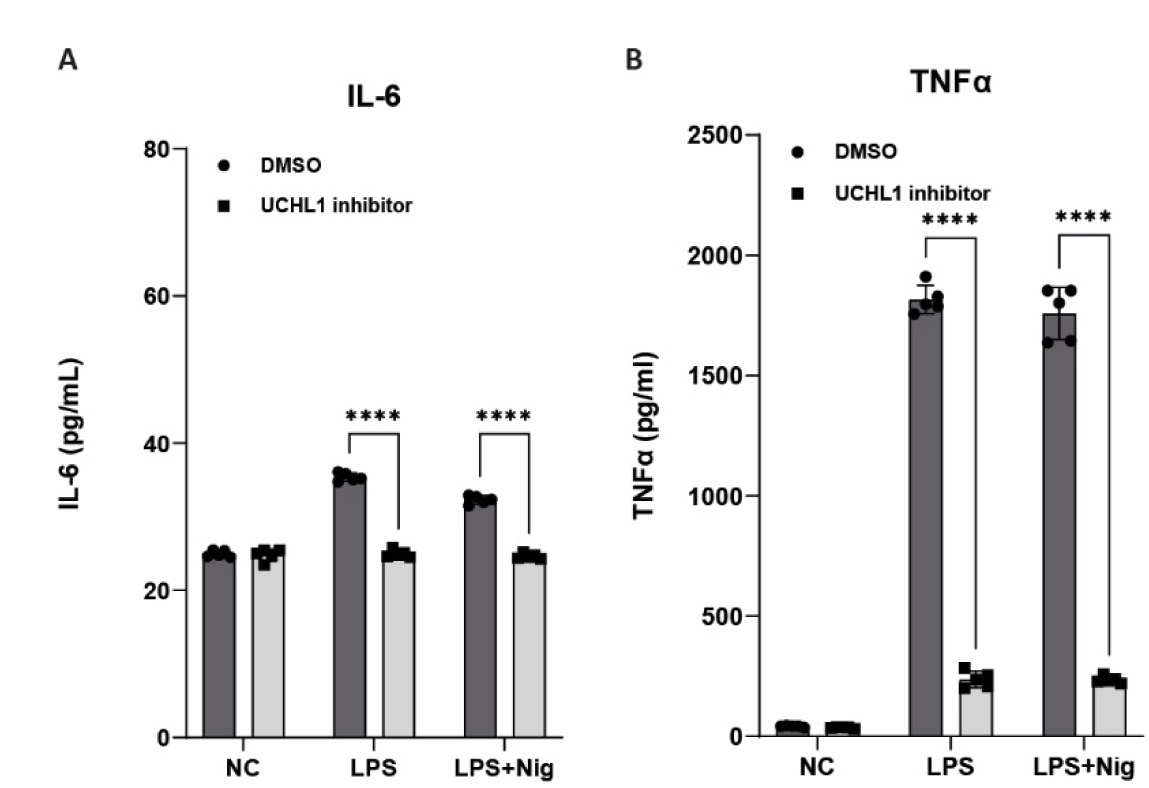
UCH-L1 inhibitor diminishes IL-6 and TNFα expression in THP-1 cells. (A, B) IL-6 (A) and TNFα (B) levels determined by ELISA from THP-1 cells treated with DMSO or 0.3 μM IMP-1711-S. Cells were differentiated with PMA and primed with LPS 1 μg/mL) for 4 hours, followed by the treatment of nigericin (10 μM) for 45 minutes. The inhibitor or the DMSO were added at the same time with LPS. Error bars show the standard deviation of the mean. ****p < 0.0001 (unpaired two-sided t-test).

**Figure S11.**
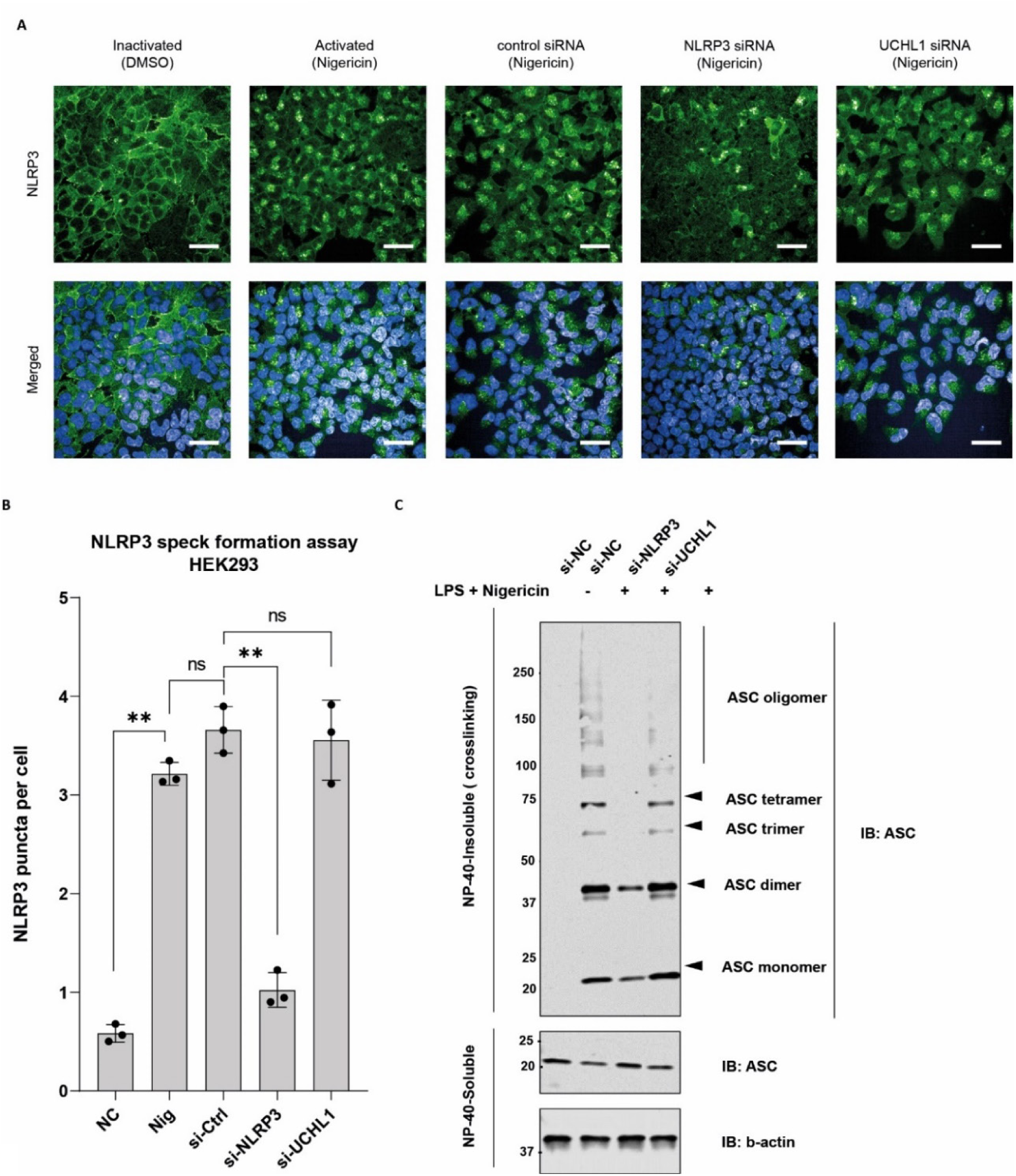
UCH-L1 is dispensable for NLRP3 puncta formation but partially alters ASC oligomerization. (A, B) flag-NLRP3 HEK293 stable cells were transfected with non-targeting control siRNA, NLRP3 or UCH-L1-specific siRNA. 48 hours after transfection, cells were treated with DMSO or 10 µM nigericin for 45 minutes. Subsequently, cells were stained with anti-flag antibody to measure the formation of NLRP3 puncta. (A) Representative fluorescence imaging of NLRP3 puncta using Opera Phenix Plus High-Content Screening System. Scale bars, 50 μm. (B) Quantitative analysis of the NLRP3 puncta formation (n = 3, mean ± s.d.). (C) Immunoblot analysis of ASC after DSS crosslinking in THP-1 cells transfected with non-targeting control siRNA, NLRP3 or UCH-L1-specific siRNA.

**Figure S12.**
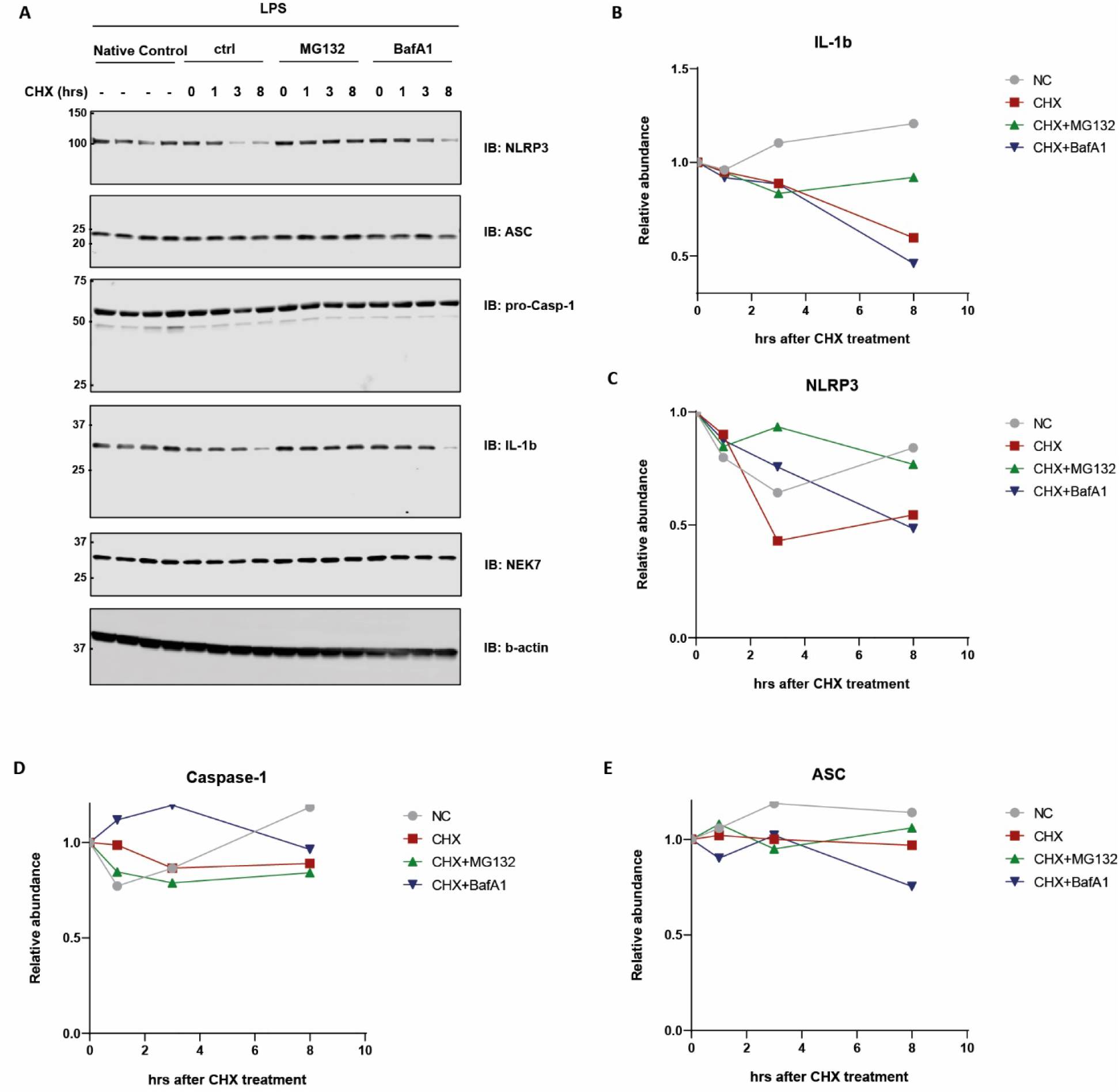
Protein turnover of NLRP3 and IL-1β depend on the ubiquitin proteasome pathway. (A) PMA-differentiated THP-1 cells were primed with LPS (100ng/mL) for 4 hours and followed by the treatment of MG132 (20μM) or BafA1 (300nM) as indicated in the last 30 min alongside. Cells were then treated with cycloheximide (50μg/mL) for 1, 3 and 8 h as indicated. Cell lysates were collected and immunoblotted for the indicated antibodies. Immunoblotting from one representative experiment of three is shown. (B-E) Quantitative analysis of the western blotting results in (A).

**Figure S13.**
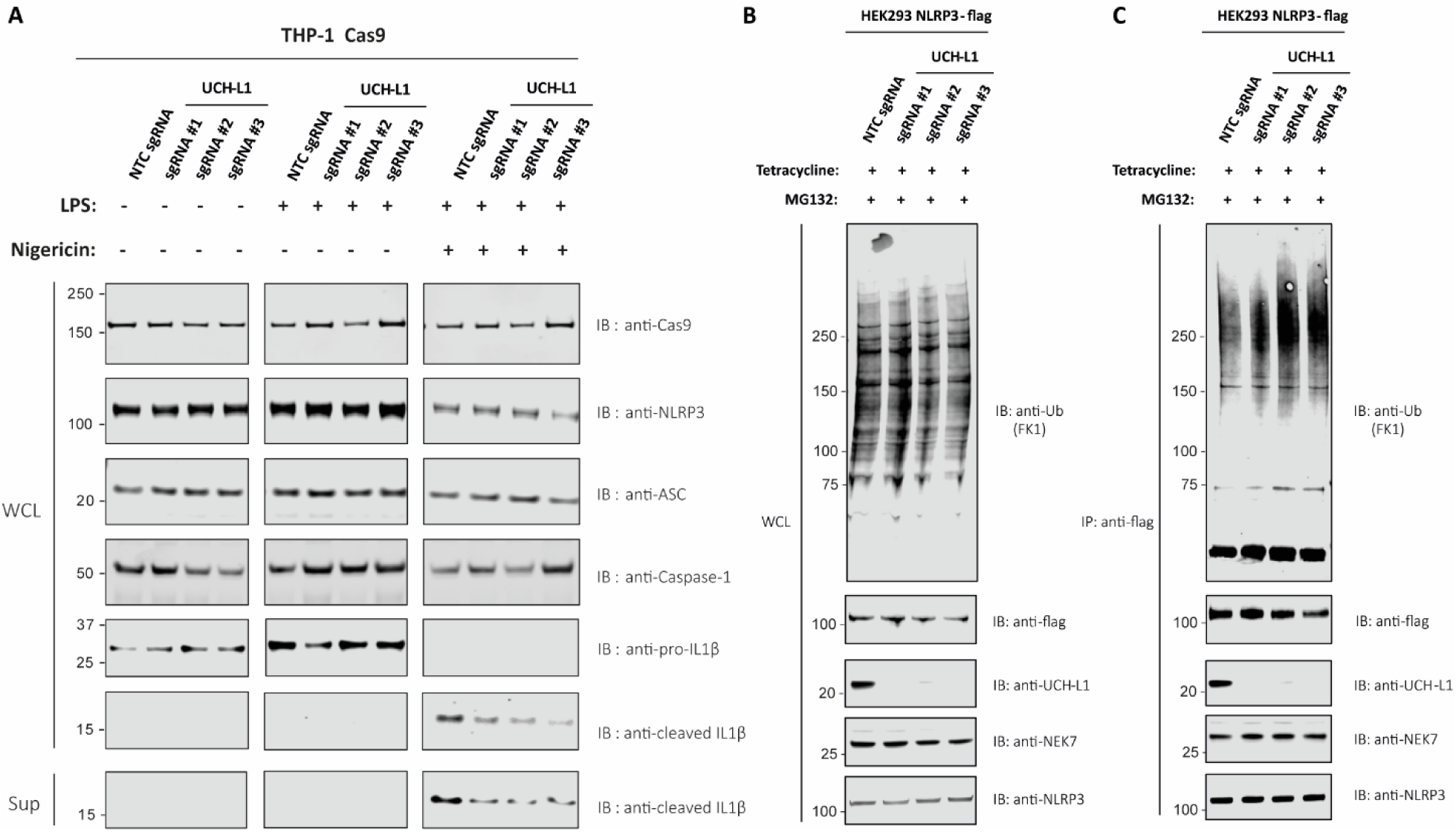
UCH-L1 deficiency abrogates IL-1β production and alters NLRP3 ubiquitination. (A) THP-1 cells, transduced by lentiviruses encoding Cas9 and non-targeting control (NTC) sgRNA or sgRNA-1/2/3 targeting UCH-L1, were differentiated with PMA, primed with LPS for 4 hours and treated with nigericin for 45 minutes, as indicated. Whole cell lysates and supernatants were immunoblotted with indicated antibodies. (B-C) flag-NLRP3 HEK293 stable cell lines, transduced by lentiviruses encoding Cas9 and non-targeting control (NTC) sgRNA or sgRNA-1/2/3 targeting UCH- L1, were treated with MG132 (10 μM) for 6 hours. Whole cell lysates (A) and flag-NLRP3 immunoprecipitated in RIPA lysis buffer (B) were analysed with indicated antibodies.

**Table S1.** List of known NLRP3-interacting proteins curated from literature.

| **Interactor** | **Experimental System** | **Throughput** | **Organism** | **Reference** |
| --- | --- | --- | --- | --- |
| **HIST1H1B** | Affinity Capture-MS | High Throughput | Homo sapiens | PUBMED:26496610 |
| **HNRNPC** | Affinity Capture-MS | High Throughput | Homo sapiens | PUBMED:26496610 |
| **RBBP7** | Affinity Capture-MS | High Throughput | Homo sapiens | PUBMED:26496610 |
| **SNRPB** | Affinity Capture-MS | High Throughput | Homo sapiens | PUBMED:26496610 |
| **SPAST** | Affinity Capture-MS | High Throughput | Homo sapiens | PUBMED:26496610 |
| **TOP1** | Affinity Capture-MS | High Throughput | Homo sapiens | PUBMED:26496610 |
| **LTBP4** | Affinity Capture-MS | High Throughput | Homo sapiens | PUBMED:26496610 |
| **SMARCA5** | Affinity Capture-MS | High Throughput | Homo sapiens | PUBMED:26496610 |
| **GEMIN2** | Affinity Capture-MS | High Throughput | Homo sapiens | PUBMED:26496610 |
| **YBX3** | Affinity Capture-MS | High Throughput | Homo sapiens | PUBMED:26496610 |
| **AP3B1** | Affinity Capture-MS | High Throughput | Homo sapiens | PUBMED:26496610 |
| **PHF14** | Affinity Capture-MS | High Throughput | Homo sapiens | PUBMED:26496610 |
| **URB1** | Affinity Capture-MS | High Throughput | Homo sapiens | PUBMED:26496610 |
| **DNAJA2** | Affinity Capture-MS | High Throughput | Homo sapiens | PUBMED:26496610 |
| **CBX3** | Affinity Capture-MS | High Throughput | Homo sapiens | PUBMED:26496610 |
| **FBXO28** | Affinity Capture-MS | High Throughput | Homo sapiens | PUBMED:26496610 |
| **CBX5** | Affinity Capture-MS | High Throughput | Homo sapiens | PUBMED:26496610 |
| **MTCH1** | Affinity Capture-MS | High Throughput | Homo sapiens | PUBMED:26496610 |
| **PVRL3** | Affinity Capture-MS | High Throughput | Homo sapiens | PUBMED:26496610 |
| **FBXL6** | Affinity Capture-MS | High Throughput | Homo sapiens | PUBMED:26496610 |
| **UQCRQ** | Affinity Capture-MS | High Throughput | Homo sapiens | PUBMED:26496610 |
| **COPS7A** | Affinity Capture-MS | High Throughput | Homo sapiens | PUBMED:26496610 |
| **LUC7L3** | Affinity Capture-MS | High Throughput | Homo sapiens | PUBMED:26496610 |
| **GPATCH4** | Affinity Capture-MS | High Throughput | Homo sapiens | PUBMED:26496610 |
| **FEM1A** | Affinity Capture-MS | High Throughput | Homo sapiens | PUBMED:26496610 |
| **LUC7L** | Affinity Capture-MS | High Throughput | Homo sapiens | PUBMED:26496610 |

| Interactor | Experimental System | Throughput | Organism | Reference |
| --- | --- | --- | --- | --- |
| **DMAP1** | Affinity Capture-MS | High Throughput | Homo sapiens | PUBMED:26496610 |
| **PHRF1** | Affinity Capture-MS | High Throughput | Homo sapiens | PUBMED:26496610 |
| **CCDC86** | Affinity Capture-MS | High Throughput | Homo sapiens | PUBMED:26496610 |
| **PCGF5** | Affinity Capture-MS | High Throughput | Homo sapiens | PUBMED:26496610 |
| **ZC3H8** | Affinity Capture-MS | High Throughput | Homo sapiens | PUBMED:26496610 |
| **KIAA2013** | Affinity Capture-MS | High Throughput | Homo sapiens | PUBMED:26496610 |
| **NLRP5** | Affinity Capture-MS | High Throughput | Homo sapiens | PUBMED:26496610 |
| **STUB1** | Affinity Capture-MS | Low Throughput | Homo sapiens | PUBMED:25594175 |
| **MARCH7** | Affinity Capture-MS | Low Throughput | Homo sapiens | PUBMED:25594175 |
| **UBR5** | Affinity Capture-MS | Low Throughput | Homo sapiens | PUBMED:25594175 |
| **CUL4A** | Affinity Capture-MS | High Throughput | Homo sapiens | PUBMED:30945288 |
| **CUL4B** | Affinity Capture-MS | High Throughput | Homo sapiens | PUBMED:30945288 |
| **VDR** | Affinity Capture-MS, Affinity Capture-Western, Reconstituted Complex | High Throughput | Homo sapiens | PUBMED:31866999 |
| **NLRC4** | Affinity Capture-Western | Low Throughput | Homo sapiens | PUBMED:15107016 |
| **SUGT1** | Affinity Capture-Western | Low Throughput | Homo sapiens | PUBMED:17435760 |
| **HSP90AA1** | Affinity Capture-Western | Low Throughput | Homo sapiens | PUBMED:17435760 |
| **DHX33** | Affinity Capture-Western | Low Throughput | Homo sapiens | PUBMED:25172487 |
| **ULK1** | Affinity Capture-Western | Low Throughput | Homo sapiens | PUBMED:26347139 |
| **MEFV** | Affinity Capture-Western | Low Throughput | Homo sapiens | PUBMED:26347139 |
| **HDAC6** | Affinity Capture-Western | Low Throughput | Homo sapiens | PUBMED:26471297 |
| **USP50** | Affinity Capture-Western | Low Throughput | Homo sapiens | PUBMED:28094437 |
| **ARIH2** | Affinity Capture-Western | Low Throughput | Homo sapiens | PUBMED:29021376 |
| **TXNIP** | Affinity Capture-Western, Two-hybrid | Low Throughput | Homo sapiens | PUBMED:28326454, PUBMED:30833078, PUBMED:32323100, PUBMED:32716108 |
| **RBX1** | Affinity Capture-Western | Low Throughput | Homo sapiens | PUBMED:30653357 |
| **FAM175B** | Affinity Capture-Western | Low Throughput | Homo sapiens | PUBMED:30787184 |
| **BRCC3** | Affinity Capture-Western | Low Throughput | Homo sapiens, Mus musculus | PUBMED:30787184, PUBMED:31866999, PUBMED:32941940 |
| **ORF8b** | Affinity Capture-Western | Low Throughput | Severe acute respiratory syndrome-related coronavirus | PUBMED:31231549 |
| **SRC** | Affinity Capture-Western | Low Throughput | Homo sapiens | PUBMED:31553908 |
| **NEK7** | Affinity Capture-Western | Low Throughput | Homo sapiens | PUBMED:31762063, PUBMED:31866999 |
| **Ppp2ca** | Affinity Capture-Western | Low Throughput | Homo sapiens | PUBMED:31866999 |
| **SQSTM1** | Affinity Capture-Western | Low Throughput | Homo sapiens | PUBMED:32003754 |
| **IL1B** | Affinity Capture-Western | Low Throughput | Homo sapiens | PUBMED:32597834 |
| **EIF3F** | Affinity Capture-Western | Low Throughput | Homo sapiens | PUBMED:33243234 |
| **USP13** | Affinity Capture-Western | Low Throughput | Homo sapiens | PUBMED:33243234 |
| **PSMB8** | Affinity Capture-Western | Low Throughput | Homo sapiens | PUBMED:32782149 |
| **PARK2** | Affinity Capture-Western | Low Throughput | Homo sapiens | PUBMED:33527027 |
| **BTRC** | Affinity Capture-Western | Low Throughput | Homo sapiens | PUBMED:33976226 |
| **TRIM24** | Affinity Capture-Western | Low Throughput | Homo sapiens | PUBMED:33724611 |
| **USP5** | Affinity Capture-Western | Low Throughput | Homo sapiens | PUBMED:34486483 |
| **TRIM28** | Affinity Capture-Western | Low Throughput | Homo sapiens | PUBMED:34373456 |
| **LYN** | Affinity Capture-Western | Low Throughput | Homo sapiens | PMID: 34699250 |
| **EIF2AK2** | Affinity Capture-Western | Low Throughput | Homo sapiens | PMID: 22801494 |
| **PRKD1/2/3** | Affinity Capture-Western | Low Throughput | Homo sapiens | PMID: 28716882 |
| **FBXL2** | Affinity Capture-Western, Biochemical Activity | Low Throughput | Homo sapiens | PUBMED:26037928 |
| **PYCARD** | Affinity Capture-Western, Co-localization | Low Throughput | Homo sapiens | PUBMED:30182366, PUBMED:32003754, PUBMED:32435622 |
| **P2RY2** | Affinity Capture-Western, Co-localization | Low Throughput | Homo sapiens | PUBMED:31553908 |
| **MAVS** | Affinity Capture-Western, Co-localization | Low Throughput | Homo sapiens | PMID: 23582325 |
| **NLRP11** | Affinity Capture-Western, Co-localization | Low Throughput | Homo sapiens | PMID: 35624206 |
| **CBLB** | Affinity Capture-Western, Reconstituted Complex | Low Throughput | Homo sapiens | PUBMED:31999304 |
| **RNF125** | Affinity Capture-Western, Reconstituted Complex | Low Throughput | Homo sapiens | PUBMED:31999304 |
| **STAMBP** | Biochemical Activity, Affinity Capture-Western | Low Throughput | Homo sapiens | PUBMED:32003754, PUBMED:33253913 |
| **MCFD2** | Co-fractionation | High Throughput | Homo sapiens | PUBMED:26344197 |
| **ARHGAP17** | Co-fractionation | High Throughput | Homo sapiens | PUBMED:26344197 |
| **SH3BP1** | Co-fractionation | High Throughput | Homo sapiens | PUBMED:26344197 |
| **CASP8** | Co-localization | Low Throughput | Homo sapiens | PUBMED:30182366 |
| **MARK4** | Co-localization | Low Throughput | Homo sapiens | PUBMED:31167564 |
| **HSPD1** | Proximity Label-MS | High Throughput | Homo sapiens | PUBMED:30021884 |
| **Rnf213** | Reconstituted Complex | High Throughput | Homo sapiens, Mus musculus | PUBMED:31999304 |
| **Trim21** | Reconstituted Complex | High Throughput | Homo sapiens, Mus musculus | PUBMED:31999304 |
| **Trim47** | Reconstituted Complex | High Throughput | Homo sapiens, Mus musculus | PUBMED:31999304 |
| **Trim14** | Reconstituted Complex | High Throughput | Homo sapiens, Mus musculus | PUBMED:31999304 |
| **ORF8** | Reconstituted Complex, Affinity Capture-Western | Low Throughput | Severe acute respiratory syndrome coronavirus 2 | DOI:10.1101/2021.12.02.470978 |
| **TRIM31** | Reconstituted Complex, Affinity Capture-Western, Biochemical Activity | Low Throughput | Homo sapiens | PUBMED:27929086 |
| **PGM1** | Two-hybrid | High Throughput | Homo sapiens | PUBMED:30653357 |
| **ATP1B3** | Two-hybrid | High Throughput | Homo sapiens | PUBMED:30653357 |
| **CUL1** | Two-hybrid, Affinity Capture-Western, Co-localization, PCA | High Throughput | Homo sapiens | PUBMED:30653357 |
| **PXN** | Two-hybrid, Affinity Capture-Western, Reconstituted Complex, Co-localization | Low Throughput | Homo sapiens | PUBMED:33243234 |
| **FAF1** | Two-hybrid, Reconstituted Complex, Affinity Capture-Western | Low Throughput | Homo sapiens | PUBMED:17046979 |

**Table S2.** List of the Golgi membrane proteins differentially captured by NLRP3-APEX2 during nigericin-induced inflammasome assembly.

| Protein IDs | Protein names | Gene names | Peptides | Unique peptides | Sequence coverage [%] |
| --- | --- | --- | --- | --- | --- |
| O14828 | Secretory carrier-associated membrane protein 3 | SCAMP3 | 11 | 11 | 42.1 |
| O94979 | Protein transport protein Sec31A | SEC31A | 26 | 26 | 23 |
| O95487 | Protein transport protein Sec24B | SEC24B | 23 | 23 | 22.7 |
| P22059 | Oxysterol-binding protein 1 | OSBP | 20 | 20 | 33 |
| P53992 | Protein transport protein Sec24C | SEC24C | 25 | 24 | 31.9 |
| P61019;Q8WUD1 | Ras-related protein Rab-2A;Ras-related protein Rab-2B | RAB2A;RAB2B | 6 | 6 | 37.7 |
| P61106 | Ras-related protein Rab-14 | RAB14 | 12 | 12 | 65.6 |
| Q00610 | Clathrin heavy chain 1 | CLTC | 57 | 57 | 39.6 |
| Q08379 | Golgin subfamily A member 2 | GOLGA2 | 22 | 22 | 28 |
| Q10567 | AP-1 complex subunit beta-1 | AP1B1 | 19 | 5 | 21.6 |
| Q12907 | Vesicular integral-membrane protein VIP36 | LMAN2 | 11 | 11 | 37.9 |
| Q14789 | Golgin subfamily B member 1 | GOLGB1 | 106 | 106 | 45.4 |
| Q15436 | Protein transport protein Sec23A | SEC23A | 20 | 17 | 35.7 |
| Q15437 | Protein transport protein Sec23B | SEC23B | 24 | 21 | 37.2 |
| Q86Y82 | Syntaxin-12 | STX12 | 8 | 8 | 37.7 |
| Q8NF37 | Lysophosphatidylcholine acyltransferase 1 | LPCAT1 | 4 | 4 | 10.7 |
| Q8TBA6 | Golgin subfamily A member 5 | GOLGA5 | 22 | 22 | 41.9 |
| Q9BXF6 | Rab11 family-interacting protein 5 | RAB11FIP5 | 12 | 12 | 32.3 |
| Q9H0U4;Q92928 | Ras-related protein Rab-1B;Putative Ras-related protein Rab-1C | RAB1B;RAB1C | 4 | 2 | 29.4 |
| Q9NRW7 | Vacuolar protein sorting-associated protein 45 | VPS45 | 11 | 11 | 19.6 |
| Q9NTJ5 | Phosphatidylinositol phosphatase SAC1 | SACM1L | 11 | 11 | 19.3 |
| Q9Y2H6 | Fibronectin type-III domain-containing protein 3A | FNDC3A | 7 | 7 | 8.3 |

**Table S3.** Human DUB siRNA screening in THP-1 cells.

| LPS |  |  |  | LPS + Nigericin |  |  |  |
| --- | --- | --- | --- | --- | --- | --- | --- |
| Rank | Gene | Normalized IL1b | z-score | Rank | Gene | Normalized IL1b | z-score |
| 1 | TNFAIP3 | 2.334 | 3.451 | 1 | STAMBP | 3.064 | 2.238 |
| 2 | UCHL3 | 2.217 | 2.720 | 2 | USP39 | 2.966 | 2.015 |
| 3 | CYLD | 2.103 | 2.013 | 3 | ZRANB1 | 2.924 | 1.917 |
| 4 | USP8 | 2.016 | 1.470 | 4 | USP17L2 | 2.899 | 1.861 |
| 5 | JOSD2 | 2.010 | 1.432 | 5 | USP38 | 2.875 | 1.806 |
| 6 | USP38 | 1.996 | 1.343 | 6 | USP47 | 2.846 | 1.740 |
| 7 | USP18 | 1.992 | 1.320 | 7 | USP25 | 2.846 | 1.739 |
| 8 | JOSD1 | 1.992 | 1.319 | 8 | UBTD1 | 2.794 | 1.620 |
| 9 | USP40 | 1.983 | 1.261 | 9 | UCK2 | 2.775 | 1.576 |
| 10 | OTUD6B | 1.977 | 1.228 | 10 | UCHL3 | 2.727 | 1.465 |
| 11 | ZRANB1 | 1.976 | 1.221 | 11 | COPS5 | 2.698 | 1.399 |
| 12 | OTUD7A | 1.967 | 1.162 | 12 | USP14 | 2.689 | 1.379 |
| 13 | OTUD7B | 1.956 | 1.098 | 13 | USP8 | 2.687 | 1.375 |
| 14 | USP33 | 1.951 | 1.066 | 14 | USP33 | 2.665 | 1.323 |
| 15 | USP46 | 1.938 | 0.984 | 15 | ATXN3 | 2.656 | 1.304 |
| 16 | USP43 | 1.923 | 0.892 | 16 | USP31 | 2.608 | 1.194 |
| 17 | STAMBPL1 | 1.914 | 0.838 | 17 | UBTD2 | 2.606 | 1.187 |
| 18 | OTUB2 | 1.902 | 0.759 | 18 | USP37 | 2.518 | 0.986 |
| 19 | USPL1 | 1.900 | 0.746 | 19 | USP43 | 2.515 | 0.981 |
| 20 | OTUD5 | 1.898 | 0.736 | 20 | OTUD1 | 2.514 | 0.976 |
| 21 | USP21 | 1.895 | 0.715 | 21 | USP34 | 2.505 | 0.957 |
| 22 | USP48 | 1.893 | 0.704 | 22 | USP5 | 2.476 | 0.890 |
| 23 | USP3 | 1.870 | 0.563 | 23 | FBXO8 | 2.452 | 0.834 |
| 24 | STAMBP | 1.867 | 0.542 | 24 | USP3 | 2.449 | 0.828 |
| 25 | USP45 | 1.863 | 0.518 | 25 | USP22 | 2.433 | 0.792 |
| 26 | USP27X | 1.863 | 0.518 | 26 | STAMBPL1 | 2.397 | 0.709 |
| 27 | USP11 | 1.860 | 0.501 | 27 | BRCC3 | 2.389 | 0.691 |
| 28 | USP13 | 1.856 | 0.472 | 28 | USP42 | 2.366 | 0.637 |
| 29 | COPS5 | 1.854 | 0.464 | 29 | USP32 | 2.357 | 0.617 |
| 30 | USP6 | 1.848 | 0.422 | 30 | USP24 | 2.340 | 0.579 |
| 31 | MYSM1 | 1.845 | 0.403 | 31 | USPL1 | 2.320 | 0.532 |
| 32 | USP50 | 1.838 | 0.361 | 32 | YOD1 | 2.315 | 0.521 |
| 33 | PAN2 | 1.832 | 0.326 | 33 | USP54 | 2.313 | 0.516 |
| 34 | SEN2 | 1.832 | 0.323 | 34 | JOSD2 | 2.293 | 0.471 |
| 35 | USP5 | 1.831 | 0.317 | 35 | UCHL5 | 2.272 | 0.422 |
| 36 | USP24 | 1.831 | 0.315 | 36 | USP30 | 2.254 | 0.381 |
| 37 | PRPF8 | 1.821 | 0.259 | 37 | USP40 | 2.237 | 0.342 |
| 38 | UFD1L | 1.816 | 0.226 | 38 | USP27X | 2.233 | 0.332 |
| 39 | USP1 | 1.814 | 0.214 | 39 | USP26 | 2.206 | 0.271 |
| 40 | VCPIP1 | 1.813 | 0.208 | 40 | PRPF8 | 2.200 | 0.258 |
| 41 | PSMD14 | 1.811 | 0.197 | 41 | UFD1L | 2.189 | 0.233 |
| 42 | UBL4A | 1.801 | 0.131 | 42 | USP35 | 2.184 | 0.220 |
| 43 | UCHL5 | 1.800 | 0.123 | 43 | OTUD4 | 2.153 | 0.150 |

| LPS |  |  |  |
| --- | --- | --- | --- |
| Rank | Gene | Normalized IL1b | z-score |
| 44 | UBTD2 | 1.796 | 0.102 |
| 45 | USP39 | 1.791 | 0.067 |
| 46 | USP29 | 1.789 | 0.059 |
| 47 | BRCC3 | 1.785 | 0.032 |
| 48 | USP28 | 1.767 | -0.077 |
| 49 | USP32 | 1.763 | -0.107 |
| 50 | USP19 | 1.754 | -0.164 |
| 51 | USP7 | 1.752 | -0.173 |
| 52 | USP41 | 1.748 | -0.200 |
| 53 | MPND | 1.745 | -0.214 |
| 54 | USP34 | 1.744 | -0.220 |
| 55 | UBTD1 | 1.738 | -0.260 |
| 56 | USP4 | 1.738 | -0.262 |
| 57 | USP10 | 1.736 | -0.273 |
| 58 | USP25 | 1.730 | -0.307 |
| 59 | USP44 | 1.729 | -0.317 |
| 60 | USP9Y | 1.729 | -0.317 |
| 61 | FBXO8 | 1.722 | -0.362 |
| 62 | USP12 | 1.721 | -0.368 |
| 63 | YOD1 | 1.721 | -0.369 |
| 64 | USP15 | 1.718 | -0.388 |
| 65 | UBL5 | 1.716 | -0.397 |
| 66 | USP22 | 1.715 | -0.405 |
| 67 | PARP11 | 1.714 | -0.408 |
| 68 | USP30 | 1.713 | -0.416 |
| 69 | USP49 | 1.710 | -0.436 |
| 70 | UEVLD | 1.710 | -0.437 |
| 71 | OTUB1 | 1.699 | -0.501 |
| 72 | USP17L2 | 1.692 | -0.545 |
| 73 | ATXN3 | 1.690 | -0.557 |
| 74 | UBL3 | 1.689 | -0.563 |
| 75 | USP47 | 1.683 | -0.603 |
| 76 | USP14 | 1.680 | -0.619 |
| 77 | USP42 | 1.680 | -0.622 |
| 78 | USP54 | 1.678 | -0.634 |
| 79 | FBXO7 | 1.672 | -0.671 |
| 80 | USP35 | 1.660 | -0.743 |
| 81 | USP16 | 1.657 | -0.765 |
| 82 | USP53 | 1.654 | -0.781 |
| 83 | USP2 | 1.654 | -0.782 |
| 84 | USP31 | 1.626 | -0.956 |
| 85 | BAP1 | 1.626 | -0.959 |
| 86 | USP51 | 1.617 | -1.017 |
| 87 | USP37 | 1.616 | -1.022 |
| 88 | OTUD1 | 1.578 | -1.258 |

| LPS + Nigericin |  |  |  |
| --- | --- | --- | --- |
| Rank | Gene | Normalized IL1b | z-score |
| 44 | USP15 | 2.146 | 0.133 |
| 45 | USP36 | 2.126 | 0.087 |
| 46 | PARP11 | 2.098 | 0.023 |
| 47 | USP18 | 2.082 | -0.014 |
| 48 | USP2 | 2.063 | -0.058 |
| 49 | PAN2 | 1.986 | -0.234 |
| 50 | UBL5 | 1.869 | -0.501 |
| 51 | USP1 | 1.862 | -0.518 |
| 52 | BAP1 | 1.826 | -0.601 |
| 53 | USP48 | 1.822 | -0.609 |
| 54 | SEN2 | 1.814 | -0.627 |
| 55 | USP9Y | 1.796 | -0.670 |
| 56 | OTUD6B | 1.796 | -0.670 |
| 57 | FBXO7 | 1.792 | -0.679 |
| 58 | USP16 | 1.785 | -0.694 |
| 59 | USP21 | 1.777 | -0.712 |
| 60 | USP29 | 1.769 | -0.733 |
| 61 | USP50 | 1.753 | -0.768 |
| 62 | UBL4A | 1.747 | -0.781 |
| 63 | USP46 | 1.741 | -0.795 |
| 64 | CYLD | 1.739 | -0.800 |
| 65 | USP28 | 1.739 | -0.801 |
| 66 | USP45 | 1.736 | -0.808 |
| 67 | OTUB2 | 1.730 | -0.820 |
| 68 | MYSM1 | 1.726 | -0.829 |
| 69 | USP49 | 1.715 | -0.855 |
| 70 | USP12 | 1.714 | -0.858 |
| 71 | UBL3 | 1.683 | -0.930 |
| 72 | USP44 | 1.679 | -0.938 |
| 73 | OTUD7A | 1.675 | -0.948 |
| 74 | TNFAIP3 | 1.668 | -0.963 |
| 75 | PSMD14 | 1.667 | -0.965 |
| 76 | USP20 | 1.664 | -0.972 |
| 77 | USP41 | 1.664 | -0.972 |
| 78 | USP4 | 1.664 | -0.973 |
| 79 | USP19 | 1.660 | -0.981 |
| 80 | MPND | 1.660 | -0.981 |
| 81 | OTUD5 | 1.658 | -0.987 |
| 82 | USP9X | 1.657 | -0.989 |
| 83 | USP53 | 1.654 | -0.996 |
| 84 | USP13 | 1.648 | -1.010 |
| 85 | JOSD1 | 1.647 | -1.011 |
| 86 | VCPIP1 | 1.640 | -1.028 |
| 87 | USP7 | 1.622 | -1.068 |
| 88 | UCHL1 | 1.617 | -1.079 |

**LPS**
| Rank | Gene | Normalized IL1b | z-score |
| --- | --- | --- | --- |
| 89 | OTUD4 | 1.577 | -1.262 |
| 90 | UBR1 | 1.520 | -1.617 |
| 91 | UCK2 | 1.513 | -1.663 |
| 92 | USP9X | 1.483 | -1.845 |
| 93 | USP20 | 1.453 | -2.034 |
| 94 | USP36 | 1.447 | -2.074 |
| 95 | UCHL1 | 1.413 | -2.286 |
| 96 | USP26 | 1.316 | -2.887 |

**LPS + Nigericin**
| Rank | Gene | Normalized IL1b | z-score |
| --- | --- | --- | --- |
| 89 | UEVLD | 1.617 | -1.081 |
| 90 | OTUB1 | 1.613 | -1.088 |
| 91 | USP11 | 1.605 | -1.108 |
| 92 | USP51 | 1.600 | -1.118 |
| 93 | OTUD7B | 1.587 | -1.149 |
| 94 | UBR1 | 1.548 | -1.238 |
| 95 | USP6 | 1.547 | -1.240 |
| 96 | USP10 | 1.539 | -1.259 |

